# BLTP3A dysfunction unleashes lysosomal stress–induced antitumor immunity through STING

**DOI:** 10.64898/2026.09.25.752224

**Authors:** Elaheh S. Hosseini, Steven Wang, Mona Arabzadeh, Kristen E. Warrington, Maria T. Søgaard, Amartya Singh, Wenjin Chen, Lomaani Ranasinghe, Rinkee Kumari, Christopher K. Alcott, Adam Robinson, Shreya Yadav, Wenwei Hu, Jian Cao, Eugenia Girda, Eileen P. White, Christian S. Hinrichs, Kyle Kristopher Payne

**Author notes:** CORRESPONDENCE: Kyle Kristopher Payne, PhD Rutgers Cancer Institute 195 Little Albany Street New Brunswick, New Jersey USA.

## Abstract

High-grade serous ovarian cancer (HGSOC) progresses within an ascites-rich environment that induces lysosomal stress. How recovery from this stress shapes protective antitumor immunity remains unclear. We identify BLTP3A as a regulator of lysosomal stress adaptation, with a common germline M1098T variant associated with improved survival and cytotoxic T cell–myeloid organization in HGSOC. STING promotes lysosomal stress-recovery through TFEB and CASM, and we uncover a STING–ATF4– BLTP3A feedback circuit that limits inflammation during this response. We find BLTP3A dysfunction compromises CASM-associated stress resolution, thereby sustaining STING signaling. In syngeneic tumors, *Bltp3a* loss promotes IFN-λ-dependent *Cxcl9*⁺ antigen-presenting macrophages to support local T cell-mediated tumor control through STING. Humanized BLTP3A^M1098T^-expressing tumors show similar macrophage remodeling, while epithelial lysosomal stress-response programs in HGSOC are associated with neighboring *CXCL9*⁺ myeloid populations. Together, these findings reveal how impaired lysosomal stress resolution sustains STING signaling to organize protective myeloid–T cell immunity in cancer.

**GRAPHICAL ABSTRACT:** 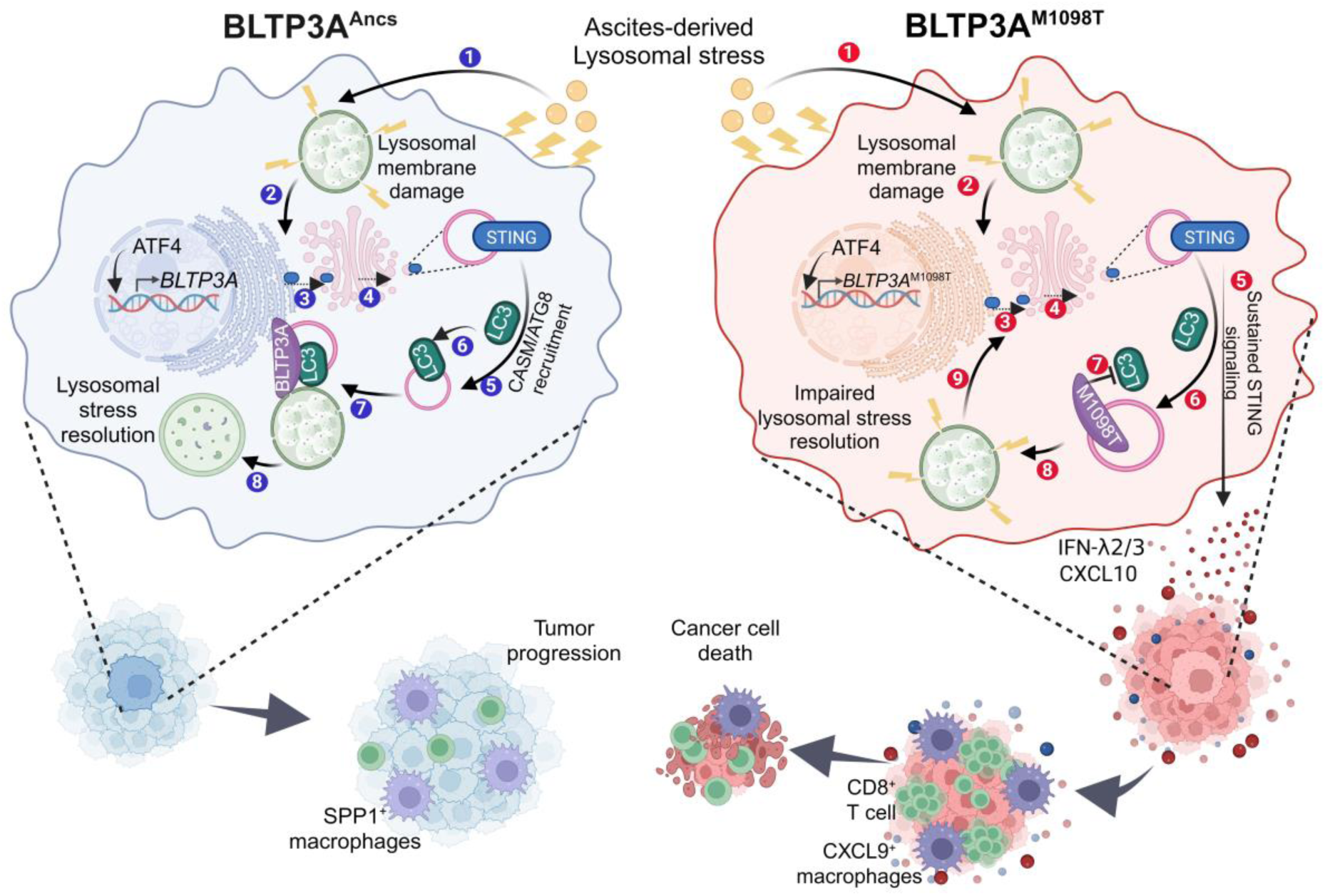

## INTRODUCTION

High-grade serous ovarian cancer (HGSOC), which accounts for 70-80% of ovarian cancer deaths, is characterized by frequent recurrence and limited durable responses to therapy^1,2^. Although HGSOC is immunogenic and frequently contains tumor-reactive lymphocytes^3–6^, productive antitumor immunity depends in part on the spatial and functional coordination of T cells with myeloid cells, with such networks associated with endogenous tumor control and responses to adoptive cellular therapy^7–10^. Tumor-derived interferons and chemokines can promote lymphocyte recruitment and shape myeloid cell states that support antitumor T cell responses^11–14^. Conversely, tumor-intrinsic programs that suppress type I interferon signaling and promote prostaglandin E₂ production can disrupt these protective interactions^7^. While these immunomodulatory signals are increasingly known, how tumor cell stress-resolution programs regulate their production to shape protective myeloid–T cell neighborhoods remain poorly understood.

Tumor cells engage stress-adaptation programs that preserve cellular fitness under adverse microenvironmental conditions. HGSOC cells disseminate through and are chronically exposed to ascites, a lipid-rich peritoneal microenvironment that imposes persistent metabolic and inflammatory stress^15–17^. Lysosomal remodeling is an important component of this adaptation, supporting nutrient recycling, pH homeostasis, drug sequestration, and membrane trafficking^18^. Extensive lysosomal membrane permeabilization can trigger cell death^19,20^, whereas sublethal injury activates membrane repair, lysophagy, lysosomal biogenesis, and ATG8-associated quality-control pathways that restore organelle integrity^21,22^. These restorative programs promote recovery from lysosomal injury and preserve tumor cell fitness. How incompletely resolved, sublethal lysosomal stress influences cancer progression, however, remains poorly understood.

Stimulator of interferon genes (STING) lies at the intersection of innate immune signaling and lysosomal homeostasis. Following activation, STING traffics from the endoplasmic reticulum to the Golgi, where it activates TBK1–IRF3-dependent interferon and inflammatory signaling, before endolysosomal processing and degradation terminate the response^15^. STING also engages lysosomal stress adaptive pathways, including TFEB-dependent lysosomal biogenesis and V-ATPase–ATG16L1-dependent conjugation of ATG8 to single membranes (CASM), which together promote recovery from lysosomal stress^21,22^. That STING participates in lysosomal stress adaptation raises an unresolved question: whether failure of the recovery program itself can convert transient, homeostatic STING activation into a persistent inflammatory state with consequences for tumor immunity.

Here, we find that HGSOC patients carrying the common germline M1098T variant in *BLTP3A*, a recently identified late-endocytic effector of CASM^23^, have longer overall survival and tumors enriched for spatially organized cytotoxic T cell–myeloid neighborhoods. We identify BLTP3A as a tumor cell- intrinsic regulator of lysosomal stress resolution and show that BLTP3A^M1098T^ impairs its stress-induced engagement with LC3-associated machinery. We determined that ovarian cancer ascites induces lysosomal stress and activates a STING–ATF4–BLTP3A feedback program, whereas BLTP3A dysfunction prolongs STING signaling and promotes IFN-λ-dependent inflammatory myeloid remodeling. This response emerges independently of adaptive immunity and is accompanied by enrichment of CXCL9⁺ antigen-presenting macrophages and the establishment of protective myeloid niches that support local CD8⁺ T cell proliferation and tumor control. In human HGSOC, *CXCL9*⁺ APC- like myeloid populations are similarly enriched near epithelial regions exhibiting elevated lysosomal remodeling. Thus, dysfunction of BLTP3A-dependent lysosomal stress-resolution programs sustains STING signaling and reshapes macrophage composition and the spatial organization of protective myeloid–T cell immunity.

## RESULTS

### A conserved germline *BLTP3A*^M1098T^ variant marks immune-mediated ovarian cancer control

Host germline variation can shape the composition of the tumor-immune microenvironment, antigen presentation, interferon signaling, and response to immunotherapy^24,25^. We therefore reasoned that inherited coding variants might reveal regulators of ovarian cancer immune control. Analysis of germline variation and clinical outcomes in the TCGA ovarian cancer cohort identified rs13205210, a missense variant in *BLTP3A* encoding the conserved M1098T substitution, as associated with improved overall survival. *BLTP3A*^M1098T^ carriers had significantly longer overall survival than patients homozygous for the ancestral allele (n = 76 carriers versus n = 243 ancestral; adjusted HR, 0.64; 95% CI, 0.45–0.91; Wald P = 0.013; Fig. 1A). We prioritized this variant for further investigation because previous genetic studies had associated *BLTP3A*^M1098T^ with susceptibility to systemic lupus erythematosus^26,27^

**Figure 1.**
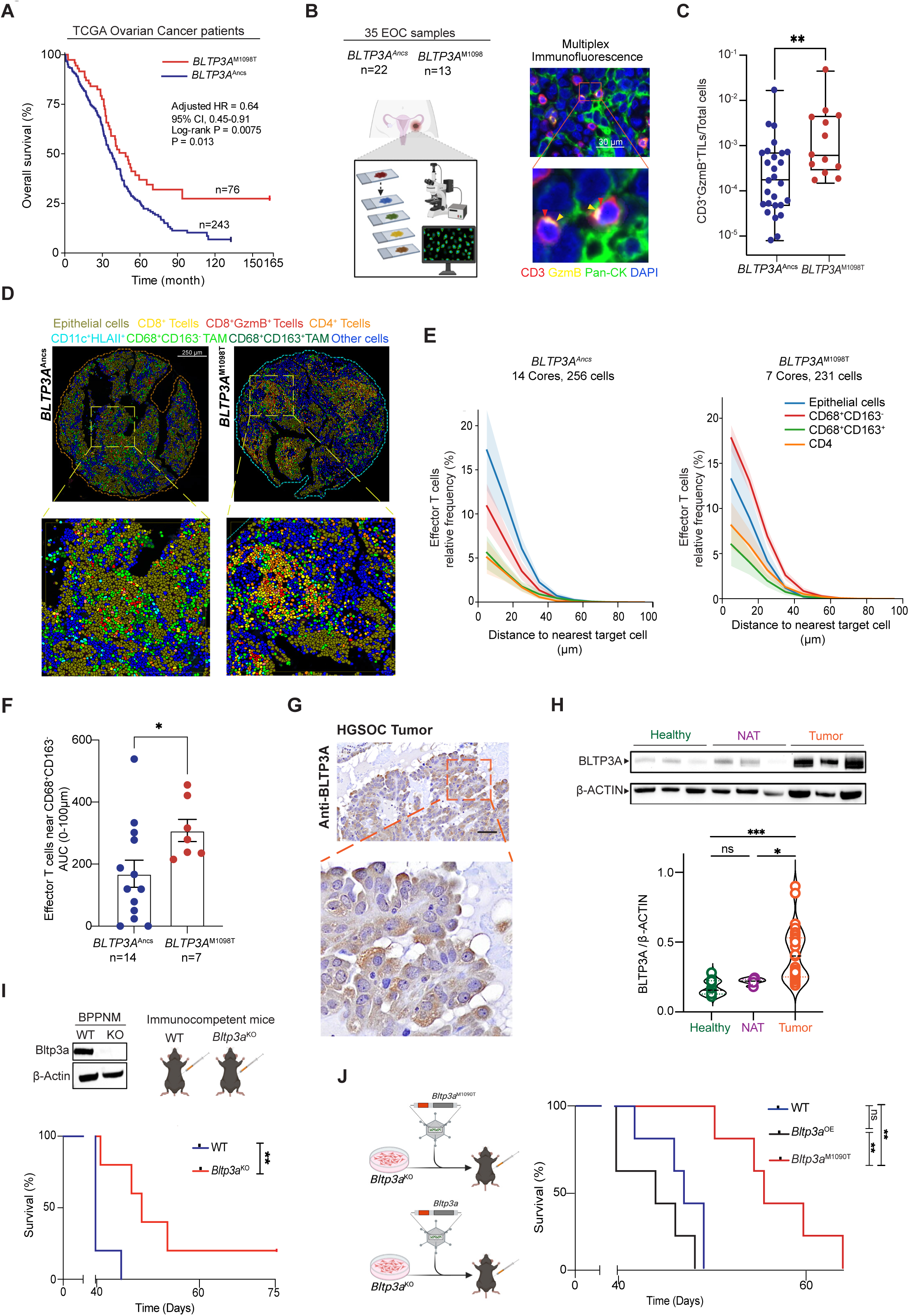
A germline BLTP3A variant associated with immune-mediated ovarian cancer control. **(A)** TCGA-OV overall survival by *BLTP3A* M1098T genotype (carriers, n = 76; ancestral, n = 243); Cox model adjusted for age, stage, and genetic ancestry; unadjusted log-rank *p* value shown. **(B)** Genotyped HGSOC specimens (ancestral, n = 22; M1098T, n = 13) and representative CD3, GzmB, pan-cytokeratin, and DAPI immunofluorescence. **(C)** CD3^+^GzmB^+^ effector T cell frequency among classified cells. **(D)** Spatial maps of epithelial, T cell, macrophage (CD68⁺CD163⁻/⁺), and CD11c⁺HLA-II⁺ myeloid populations. **(E)** CD8^+^GzmB^+^ T cell relative frequency by distance to the indicated cell populations. **(F)** Area under 0–100-μm proximity curves for CD8⁺GzmB⁺ T cells near CD68⁺CD163⁻ macrophages. **(G)** Representative BLTP3A immunohistochemistry in human HGSOC. **(H)** BLTP3A immunoblot and normalized quantification in healthy (n = 9), normal-adjacent (n = 4), and HGSOC (n = 18) tissues. **(I)** *Bltp3a* knockout validation and survival following intraperitoneal WT or KO BPPNM challenge. **(J)** Survival following WT challenge or *Bltp3a*^KO^ challenge with ancestral BLTP3A or orthologous M1090T reconstitution. Data are mean ± SEM unless otherwise indicated; each point represents an independent tumor or biological replicate. Survival curves were compared by log-rank tests; other comparisons used the tests described in Methods. \**p* < 0.05, \*\**p* < 0.01, \*\*\**p* < 0.001; ns, not significant.

This association with survival and autoimmune susceptibility prompted us to determine whether *BLTP3A*^M1098T^ carrier tumors also exhibited increased intratumoral T cell activity. We genotyped HGSOCs from our tumor bank and performed multiplex immunofluorescence on 35 FFPE tumors, including 22 tumors homozygous for the ancestral allele and 13 tumors carrying *BLTP3A*^M1098T^ in heterozygosity or homozygosity (Fig. 1B). Compared with ancestral tumors, *BLTP3A*^M1098T^ carrier tumors were significantly enriched for CD3^+^GzmB^+^ effector T cells (Fig. 1C), which frequently localized within tumor epithelial islets (Fig. S1A), consistent with tumor-proximal cytotoxic immune activity^3,4,9,13^. Further stratification by genotype showed that effector T cell enrichment was greatest in tumors homozygous for *BLTP3A*^M1098T^ (Fig. S1B).

Because spatial immune organization is a defining feature of productive antitumor immunity^4,9,11^, we performed sequential immunofluorescence (seqIF), cellular phenotyping and distance-based nearest- neighbor analysis in a genotyped subset of HGSOCs comprising 14 tumors homozygous for the ancestral allele and 7 carrying *BLTP3A*^M1098T^ (Fig. 1D). seqIF confirmed increased T cell frequencies in *BLTP3A*^M1098T^ carrier tumors and further revealed greater intratumoral myeloid cell abundance (Fig. S1C, D). Nearest-neighbor analysis revealed increased proximity between CD8⁺GzmB⁺ effector T cells and CD68⁺CD163⁻ tumor-associated macrophages (TAMs) in *BLTP3A*^M1098T^ carrier tumors reflected by a greater area under the proximity curve over distance of 0-100 μm (Fig. 1E-F). Thus, *BLTP3A*^M1098T^ is associated with an immune-rich spatial phenotype characterized by closer coupling of effector T cells with macrophages lacking the canonical M2-associated marker CD163, potentially supporting local antitumor myeloid–T cell interactions.

Since rs13205210 is germline encoded, we next examined the cellular distribution of BLTP3A in HGSOC tissues to identify cellular compartments potentially impacted by its biological effects. Immunostaining revealed pronounced cytoplasmic BLTP3A expression in malignant epithelial cells, with comparatively little signal in adjacent stroma (Fig. 1G). Consistent with this localization, immunoblotting of healthy tissue, normal adjacent tissue (NAT), and tumor specimens showed enrichment of BLTP3A protein in tumors (Fig. 1H). Analysis of an integrated single-cell RNA-sequencing atlas comprising 505,102 cells from 84 patients with ovarian cancer further demonstrated increased *BLTP3A* expression in malignant epithelial cells relative to nonmalignant adjacent and benign epithelial populations^28^ (Fig. S1E). Together, these analyses identified malignant epithelial cells as a prominent BLTP3A-expressing compartment in HGSOC. Notably, *BLTP3A*^M1098T^ carrier tumors retained detectable BLTP3A protein levels comparable to ancestral tumors (Fig. S1F), suggesting that the M1098T variant is not associated with markedly reduced BLTP3A protein abundance.

### Tumor cell-intrinsic Bltp3a restrains immune-mediated ovarian cancer control

To determine whether tumor cell-intrinsic Bltp3a activity restrains immune control of ovarian cancer *in vivo*, we first ablated *Bltp3a* in the syngeneic BPPNM model of HGSOC^29^. CRISPR/Cas9-mediated *Bltp3a* deletion significantly prolonged survival in immunocompetent hosts relative to wild-type (WT) controls (Fig. 1I). Longitudinal bioluminescence imaging further demonstrated sustained suppression of tumor progression in mice bearing *Bltp3a*^KO^ tumors, accompanied by reduced endpoint tumor burden (Fig. S1G, H). Approximately 10% of mice challenged with *Bltp3a*^KO^ BPPNM cells rejected their tumors and subsequently resisted rechallenge with WT BPPNM cells, consistent with the establishment of protective antitumor immune memory (Fig. S1I). In contrast, *Bltp3a* loss conferred no survival advantage in NSG mice, demonstrating that the protective effect of tumor-cell-intrinsic *Bltp3a* deficiency requires host immunity (Fig. S1J). Conditional *Bltp3a* ablation similarly prolonged survival in an inducible Kras-driven, p53-deficient orthotopic ovarian cancer model (Fig. S1K).

We next tested whether the orthologous M1098T substitution recapitulated the effect of *Bltp3a* loss *in vivo*. M1098 resides within a highly conserved disordered region of BLTP3A, with murine M1090 representing the orthologous residue (Fig. S1L). Reconstitution of *Bltp3a^KO^* BPPNM cells with WT *Bltp3a* (*Bltp3a*^OE^) restored aggressive tumor growth, whereas *Bltp3a^M1090T^* expression also significantly delayed malignant progression (Fig. 1J). These findings establish M1090T as a loss-of-function-like variant *in vivo* and identify tumor-cell-intrinsic BLTP3A as a regulator of ovarian cancer progression.

### *Bltp3a* loss generates spatially organized protective CD8^+^ T cell immunity

The increased spatial proximity between cytotoxic T cells and macrophages observed in HGSOCs carrying *BLTP3A* prompted us to investigate whether tumor cell-intrinsic *Bltp3a* loss influences the composition and functional states of the tumor-immune infiltrate. We therefore performed single-cell RNA sequencing (scRNA-seq) of intratumoral CD45^+^ leukocytes isolated from mice bearing syngeneic WT and *Bltp3a*^KO^ intraperitoneal BPPNM tumors. Immune populations were annotated using canonical lineage and state-defining gene-expression programs (Fig. S2A). Genotype-separated UMAPs revealed broad remodeling of the immune landscape, with WT tumors dominated by myeloid populations and *Bltp3a*^KO^ tumors showed a prominent expansion of T cells (Fig. 2A). Quantification of the annotated subsets revealed marked remodeling of macrophage and T cell compartments. WT tumors were enriched for *Spp1*⁺ and *C1qc*⁺ macrophages alongside exhausted CD8⁺ T cells, whereas *Bltp3a*^KO^ tumors exhibited enrichment of *Cxcl9*⁺ TAMs and expansion of effector and effector-memory CD8⁺ T cell populations (Fig. 2B). This shift toward *Cxcl9*⁺ macrophages is notable given the prognostic significance of CXCL9–SPP1 macrophage polarity across cancers, including ovarian cancer^30^. CXCL9-producing macrophages support antitumor immunity through CD8⁺ T cell recruitment and antigen presenting capacities^31^.

**Figure 2.**
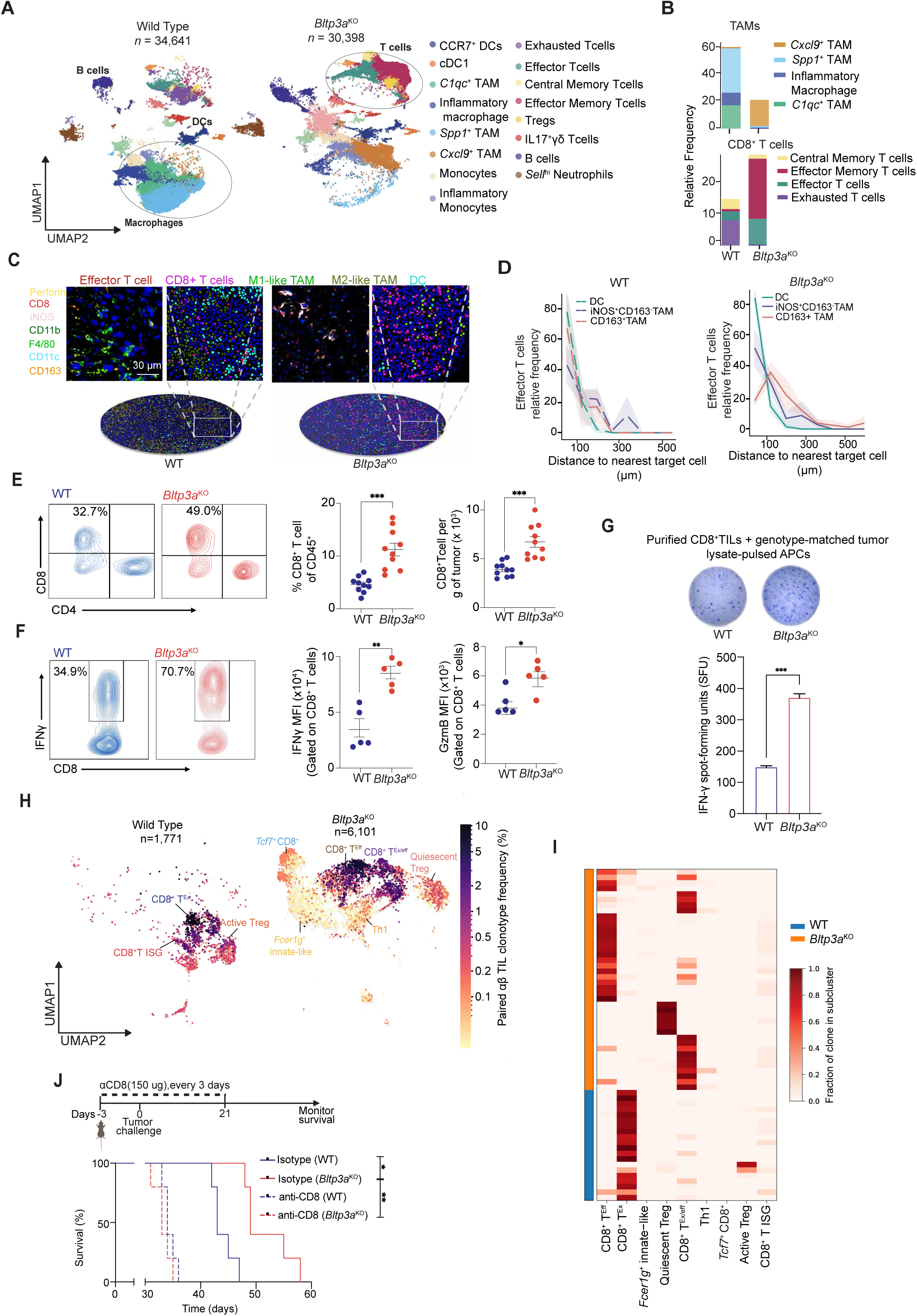
Loss of *Bltp3a* in tumor cells generates spatially organized protective CD8^+^ T cell immunity. **(A)** Single-cell RNA-seq UMAPs of intratumoral CD45⁺ leukocytes (WT, n = 34,641; KO, n = 30,398), annotated by immune-cell state. **(B)** Frequencies of TAM and CD8⁺ T-cell subsets. **(C)** Representative multiplex immunofluorescence maps identifying effector and total CD8⁺ T cells, iNOS⁺CD163⁻ and CD163⁺ macrophages, and CD11c^+^ dendritic cells. **(D)** Effector T cell frequency by distance to indicated myeloid populations. **(E)** CD8⁺ TIL frequency and absolute number per gram of tumor. (**F**) CD8⁺ TIL IFNγ and granzyme B expression following *ex vivo* stimulation. (**G**) IFNγ ELISPOT of purified CD8⁺ TILs cocultured with antigen-presenting cells pulsed with WT or KO tumor lysate. (**H**) Productive paired TCRαβ clonotypes mapped onto T cell UMAPs, colored by clonotype frequency. (**I**) Clonotype distribution across T cell subclusters. (**J**) CD8-depletion schedule and tumor-bearing mouse survival (n = 5/group). Data are mean ± SEM unless otherwise indicated; each point represents an independent tumor or biological replicate. Survival curves were compared by log-rank tests; other comparisons used the tests described in Methods. \**p* < 0.05, \*\**p* < 0.01, \*\*\**p* < 0.001; ns, not significant.

To further characterize the T cell responses associated with *Bltp3a* loss, we subclustered intratumoral T cells into transcriptional states encompassing effector-associated (*Ccl5, Nkg7 and Gzmk)*, exhaustion- associated (*Pdcd1, Lag3 and Tigit)*, and naive/memory-associated (*Tcf7, Ccr7 and Lef1)* (Figs. S2B, C). Compared with WT tumors, *Bltp3a*^KO^ tumors were enriched for effector and *Tcf7*-associated CD8⁺ T cell populations (Fig. S2D). Feature plots of *Gzmk*, *Ctla4*, and *Sell* further supported the effector, activated- Treg, and *Tcf7/Sell*-associated cluster annotations, respectively (Fig. S2E). Additional changes included increased *Fcer1g*⁺ innate-like T cells, quiescent Tregs, and A20⁺ CD4⁺ T cells. CD8⁺ T cells in *Bltp3a*^KO^ tumors also exhibited increased expression of GITR (*Tnfrsf18*) and OX40 (*Tnfrsf4*) alongside inhibitory receptors (Fig. S2F), further supporting an activated effector/exhausted-like transcriptional state^32–34^. Together, these findings suggest that loss of *Bltp3a* in tumor cells favors an immune microenvironment supportive of antitumor CD8⁺ T cell responses.

The coordinated enrichment of *Cxcl9*⁺ macrophages and activated CD8⁺ T cells offered a potential explanation for the closer spatial association between cytotoxic T cells and CD163⁻ macrophages observed in human *BLTP3A*^M1098T^ tumors (Fig. 1E, F). We therefore asked whether the transcriptional remodeling following *Bltp3a* loss was accompanied by corresponding changes in spatial immune organization. Using seqIF (Fig. 2C), we found that T cells in WT tumors were frequently located within 100μm of M2-like (CD163⁺) macrophages, whereas *Bltp3a*^KO^ tumors exhibited localized neighborhoods enriched for effector T cells and M1-like (iNOS⁺CD163⁻) macrophages, with significantly greater spatial proximity between these populations (Figs. 2D and S2G). This reorganization was accompanied by increased intratumoral CD8⁺ T cell abundance and greater IFNγ and granzyme B production following *ex vivo* stimulation (Figs. 2E, F, S2H). Moreover, CD8⁺ tumor-infiltrating lymphocytes (TILs) from *Bltp3a*^KO^ tumors exhibited a higher frequency of IFNγ-secreting cells following stimulation with matched tumor lysates (Fig. 2G), consistent with enhanced tumor-reactive T-cell activity. Together, these findings suggest that loss of *Bltp3a* in tumor cells promotes inflammatory macrophage-associated T cell neighborhoods accompanied by heightened CD8⁺ T cell function.

We next asked whether enhanced CD8⁺ T cell activity was accompanied by changes in TCR repertoire organization. GLIPH2 analysis, which groups TCRs based on shared CDR3 sequence motifs and global similarity to identify potential shared antigen specificities^35^, revealed 126 CDR3β motifs unique to *Bltp3a*^KO^ tumors, compared with 14 unique to WT tumors and 124 shared between genotypes (Fig. S2I). Repertoire diversity also trended higher in *Bltp3a*^KO^ tumors by inverse Simpson analysis (Fig. S2J). Mapping paired TCRαβ clonotypes onto transcriptional states further revealed that expanded clonotypes in WT tumors were concentrated within the exhausted CD8⁺ compartment, whereas those in *Bltp3a*^KO^ tumors were more broadly distributed across effector and effector/exhausted-like CD8⁺ states (Figs. 2H, I). Together, these findings indicate that *Bltp3a* loss is associated with altered TCR sequence convergence and broader representation of expanded clonotypes within effector-associated CD8⁺ T cell populations.

Finally, to determine whether CD8⁺ T cells were required for improved tumor control following *Bltp3a* loss, we depleted CD8⁺ T cells in mice bearing *Bltp3a*^KO^ tumors. Depletion abolished the survival advantage conferred by *Bltp3a* loss (Figs. 2J, S2K). These findings establish that the antitumor benefit of *Bltp3a* loss requires CD8⁺ T cells and support a model in which the *Cxcl9*⁺ macrophage-enriched microenvironment favors the accumulation and functional activity of effector CD8⁺ T cells, including clonally expanded populations.

### Cancer cell-intrinsic *Bltp3a* loss establishes IFN-responsive myeloid niches sustaining local CD8⁺ T cell immunity

Having identified distinct tumor-associated myeloid states, we next asked how *Bltp3a* loss altered their functional programs. *Cxcl9*⁺ TAMs enriched in *Bltp3a*^KO^ tumors expressed genes associated with antigen processing and presentation, interferon responsiveness, and inflammatory chemokine production, including *Cxcl9*, *Cxcl10*, and *Ccl5* (Fig. 3A). Conventional type 1 dendritic cells (cDC1s) and *CCR7*^+^ DCs displayed related immunostimulatory programs. Pathway enrichment analysis further distinguished *Cxcl9*⁺ TAMs and cDC1s, which exhibited antiviral and IFNγ-response signatures, from *Spp1*⁺ and *C1q*⁺ TAMs, which enriched for hypoxia, glycolysis, and EMT-associated programs (Fig. 3B). Monocytes exhibited a distinct inflammatory profile characterized by IL-6–JAK–STAT3 signaling and responses to IFNγ and hypoxia.

**Figure 3.**
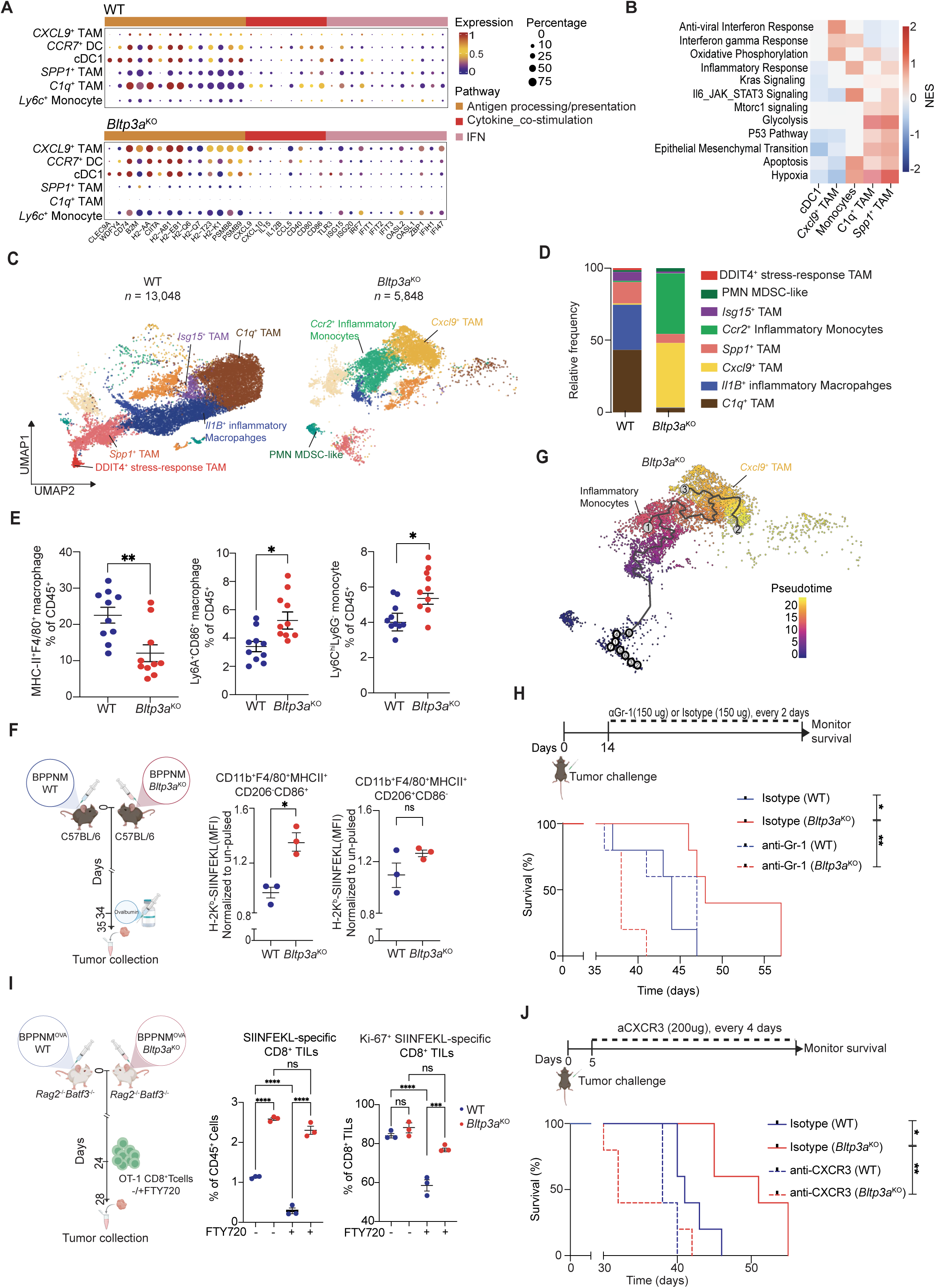
IFN-responsive myeloid populations support local CD8^+^ T cell immunity. **(A)** Myeloid cell expression of antigen-presentation, costimulatory, cytokine, and interferon-response genes. **(B)** Heatmap of normalized enrichment scores (NESs) for selected Hallmark pathways across the indicated dendritic-cell, macrophage, and monocyte states. **(C)** Genotype-separated macrophage/monocyte UMAPs showing annotated populations. **(D)** Relative frequencies of myeloid cell states. **(E)** Flow-cytometric frequencies of MHC-II⁺F4/80⁺ and Ly6A⁺CD86⁺ macrophages and Ly6C^hi^ Ly6G⁻ monocytes. **(F)** Ovalbumin-pulse assay and SIINFEKL–H-2Kᵇ presentation on CD86⁺ versus CD206⁺ TAMs, normalized to unpulsed controls. (**G**) Inferred pseudotime trajectory from *Ccr2*⁺ inflammatory monocytes toward *Cxcl9*⁺ TAMs. (**H**) Anti-Gr-1 treatment schedule and survival in WT or *Bltp3a*^KO^ BPPNM challenged mice (n=5/group). (**I**) OT-I transfer into *Rag2*⁻/⁻*Batf3*⁻/⁻ mice bearing OVA- expressing tumors, with or without FTY720 (n=3/group); SIINFEKL-specific TIL frequency and Ki67 expression measured by flow cytometry. (**J**) Anti-CXCR3 treatment schedule and survival in WT or *Bltp3a*^KO^ BPPNM challenged mice (n=5/group). Data are mean ± SEM unless otherwise indicated; each point represents an independent tumor or biological replicate. Survival curves were compared by log-rank tests; other comparisons used the tests described in Methods. \**p* < 0.05, \*\**p* < 0.01, \*\*\**p* < 0.001; ns, not significant.

To further resolve this myeloid remodeling, we subclustered macrophage and monocyte-lineage cells confirming contraction of tissue-remodeling TAM populations alongside expansion of *Cxcl9*⁺ TAMs and inflammatory myeloid populations in *Bltp3a*^KO^ tumors (Figs. 3C, D, S3A, B). Flow cytometry independently demonstrated increased Ly6C^hi^ inflammatory monocytes and IFN-responsive Ly6A⁺CD86⁺ macrophages, consistent with an expanded inflammatory myeloid compartment exhibiting IFN-associated and costimulatory features, despite reduced mature macrophage abundance (Figs. 3E, S3C). Together, these findings suggest that *Bltp3a* loss shifts the tumor myeloid environment away from metabolically adapted, tissue-remodeling programs toward interferon-responsive states associated with immune surveillance. The accompanying enrichment of antigen-presentation-associated programs prompted us to investigate whether these populations exhibited enhanced antigen-handling capabilities.

Following acute intraperitoneal administration of ovalbumin to mice bearing WT or *Bltp3a*^KO^ tumors, CD86⁺ TAMs from *Bltp3a*^KO^ tumors displayed significantly greater SIINFEKL–H-2Kᵇ presentation than their WT counterparts, whereas CD206⁺ TAMs showed no corresponding increase (Fig. 3F). These findings suggest that the *Bltp3a*^KO^ tumor microenvironment selectively enhances antigen presentation by CD86⁺ macrophages. Macrophages from *Bltp3a*^KO^ tumors also contain greater amounts of tumor-derived GFP, consistent with increased acquisition of tumor material *in vivo* (Fig. S3D). Furthermore, given recent evidence that inflammatory myeloid cells can support CD8⁺ T cell activation through acquisition of tumor- derived MHC-I (cross-dressing)^7,14^, we investigated whether this process was enhanced following *Bltp3a* loss. H-2Kᵇ-expressing *Bltp3a*^KO^ tumors were established in H-2Kᵈ BALB/c hosts under T cell-depleting conditions. CD86⁺ TAMs from *Bltp3a*^KO^ tumors exhibited greater acquisition of tumor-derived H-2Kᵇ than CD206^+^ TAMs (Fig. S3E), consistent with enhanced MHC-I cross-dressing. Together, these findings reveal enhanced antigen acquisition and presentation-associated capabilities among macrophages in *Bltp3a*^KO^ tumors.

Pseudotime analysis identified a continuous trajectory connecting *Ccr2*⁺ inflammatory monocytes with *Cxcl9*⁺ TAMs, suggesting a potential monocyte-to-macrophage differentiation axis (Fig. 3G). To investigate the contribution of inflammatory monocytes to this response, we treated mice bearing WT or *Bltp3a*^KO^ tumors with the Gr-1-depleting antibody RB6-8C5, which targets Ly6C^hi^ inflammatory monocytes and other Gr-1⁺ myeloid populations, beginning 14 days after tumor implantation. Treatment markedly reduced intratumoral Gr-1⁺ myeloid cells and F4/80⁺ macrophages (Fig. S3F) and abolished the survival advantage conferred by *Bltp3a* loss (Fig. 3H). Gr-1 depletion also eliminated the increased accumulation of CD8⁺ TILs in *Bltp3a*^KO^ tumors while having little effect on WT tumors (Fig. S3G). Thus, the enhanced CD8⁺ T cell accumulation and tumor control associated with *Bltp3a* loss depend on a Gr-1- sensitive myeloid compartment.

The loss of CD8⁺ T cell accumulation following Gr-1 depletion implicated inflammatory myeloid populations in supporting the protective T cell response. Given the established role of cDC1s in sustaining intratumoral CD8⁺ T cell responses^36^ and evidence that myeloid antigen-presenting niches can provide local T cell support through co-stimulation^9^, we asked whether *Bltp3a*^KO^ tumors could maintain antigen- specific CD8⁺ T cells despite the absence of Batf3-dependent cDC1s and restricted lymphocyte egress. *Rag*2^⁻/⁻^*Batf3*^⁻/⁻^ mice bearing ovalbumin-expressing WT or *Bltp3a*^KO^ tumors received OT-I CD8⁺ T cells with or without FTY720 to restrict lymphocyte egress. Even in hosts lacking Batf3-dependent cDC1s, *Bltp3a*^KO^ tumors contained higher frequencies of SIINFEKL-specific CD8⁺ TILs and Ki-67⁺ antigen- specific T cells than WT tumors. Importantly, FTY720 significantly reduced both populations in WT tumors but had no significant effect in *Bltp3a*^KO^ tumors (Figs. 3I, S3H). These findings suggest that *Bltp3a* loss establishes a tumor microenvironment capable of sustaining local antigen-specific CD8⁺ T cell accumulation and proliferative activity despite restricted lymphocyte egress and the absence of Batf3- dependent cDC1s, potentially through the expanded inflammatory monocyte/macrophage compartment.

The maintenance of antigen-specific CD8⁺ T cells in *Bltp3a*^KO^ tumors despite restricted lymphocyte egress suggested that local signals contribute to their accumulation and proliferative activity. Given the enrichment of *Cxcl9*⁺ TAMs expressing the CXCR3 ligands *Cxcl9* and *Cxcl10*, we investigated whether CXCR3 signaling was required for this protective response. We found that CXCR3 blockade abolished the survival advantage conferred by *Bltp3a* loss (Fig. 3J). Together, these findings support a model in which *Bltp3a* loss establishes an IFN-responsive myeloid environment that supports local antigen-specific CD8⁺ T cell immunity, with CXCR3 signaling essential for tumor control^7,14^.

### BLTP3A dysfunction unleashes tumor-derived IFN-λ-dependent antitumor immunity

Having established that tumor cell-intrinsic *Bltp3a* loss generates an IFN-responsive myeloid niche supporting protective CD8^+^ T cell immunity, we next sought to identify the tumor-derived signals that instruct this myeloid state. Because lymphocyte-derived signals, particularly IFNγ, can reinforce inflammatory and antigen-presenting macrophage programs^13,30^, we examined whether the myeloid remodeling associated with *Bltp3a* loss required adaptive immunity. In *Rag2*^-/-^ mice, *Bltp3a*^KO^ tumors continued to preferentially accumulate IFN-responsive Ly6A⁺CD86⁺ macrophages and inflammatory monocytes (Fig. 4A, S4A), demonstrating that this myeloid response can arise independently of adaptive immunity.

**Figure 4.**
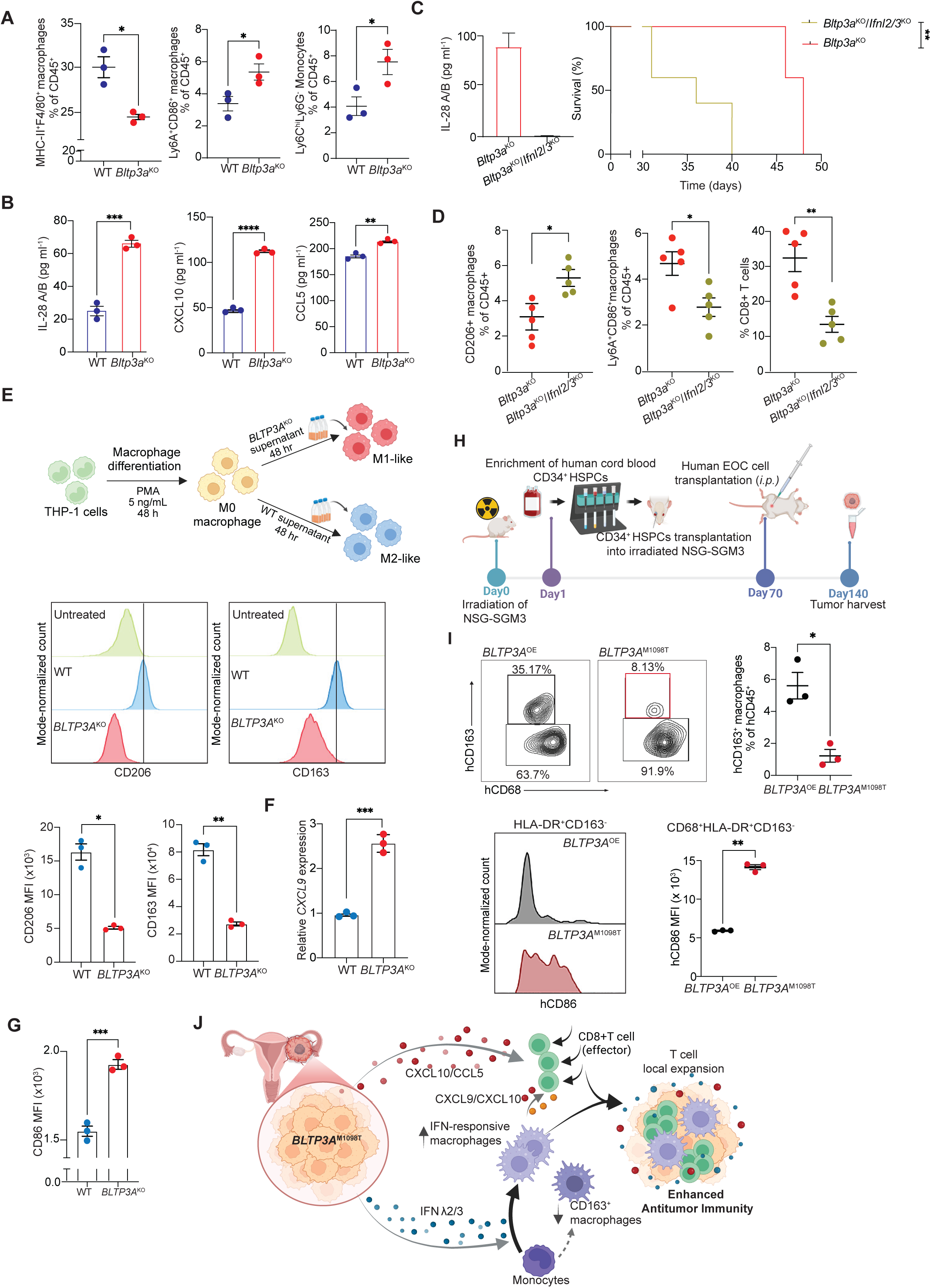
BLTP3A loss promotes tumor-derived IFN-λ-dependent immunity. **(A)** Flow-cytometric quantification of MHC-II^+^F4/80^+^ macrophages, Ly6A^+^CD86^+^ macrophages, and Ly6C^hi^Ly6G^−^ inflammatory monocytes from *Rag2*^−/−^ mice. **(B)** IL-28A/B, CXCL10 and CCL5 concentrations in conditioned media from WT and *Bltp3a*^KO^ BPPNM cells. **(C)** IFN-λ secretion following *Ifnl2/3* deletion in *Bltp3a*^KO^ cells and corresponding mouse survival. **(D)** Flow-cytometric quantification of CD206^+^ macrophages, Ly6A^+^CD86^+^ macrophages and CD8^+^ T cells in *Bltp3a*^KO^ and *Bltp3a*^KO^/*Ifnl2/3*^KO^ BPPNM tumors. (**E-G**) THP-1-derived macrophages exposed to conditioned media from WT or BLTP3A-deficient OVCAR8 cells: (E) flow-cytometric defined CD206/CD163 expression, (F) relative *CXCL9* mRNA expression, and (G) flow-cytometric defined CD86 expression. **(H)** Cord-blood-humanized NSG-SGM3 intraperitoneal OVCAR3 model. (**I**) Flow-cytometric quantification of human CD163⁺ macrophages and CD86 expression among CD68⁺HLA-DR⁺CD163⁻ macrophages from ancestral BLTP3A or M1098T- expressing OVCAR3 tumors. (**J**) Proposed model of tumor-derived IFN-λ/chemokine signaling, myeloid remodeling, and CD8⁺ T cell immunity in *BLTP3A*^M1098T^ carrying tumors. Statistical analysis was performed as follows: for (B), (D), (E), (F), and (G), two-tailed Student’s tests; *P < 0.05; **P ≤0.01; ***P ≤ 0.001.

The persistence of this myeloid remodeling suggested that an inflammatory signal independent of adaptive immunity initiates the response. Given BLTP3A’s role in endolysosomal stress adaptation, we next investigated whether its loss altered the inflammatory secretome of tumor cells. We determined that *Bltp3a*^KO^ BPPNM cells secreted increased amounts of the interferon-inducible chemokines CXCL10 and CCL5, together with IL-28A/B, the murine type III interferon IFN-λ2/3 (Fig. 4B). This was particularly notable given the prominent interferon-responsive program exhibited by *Cxcl9*⁺ macrophages enriched in *Bltp3a*^KO^ tumors (Figs. 3A, B). CRISPR/Cas9-mediated deletion of *Ifnl2/3* abolished the survival advantage conferred by *Bltp3a* loss (Fig. 4C), reduced CD8⁺ T cell accumulation, and reversed the associated macrophage remodeling, increasing M2-like CD206⁺ macrophages while reducing IFN- responsive Ly6A⁺CD86⁺ and MHC-II⁺ macrophage populations (Figs. 4D, S4B). Given the established role of type I interferons in antitumor immunity^37^, we also examined their contribution to this response. In contrast to IFN-λ, IFNAR1 blockade did not diminish the survival benefit associated with *Bltp3a* loss (Fig. S4C). Together, these findings establish that protective myeloid–T cell immunity following *Bltp3a* loss requires type III, but not type I, interferon signaling.

Having established that tumor-derived IFN-λ is required for the survival benefit conferred by *Bltp3a* loss, we next investigated its therapeutic potential. We engineered a DNA-encoded, tumor-targeted IFN-λ procytokine incorporating matrix metalloproteinase-cleavable elements and an FSH receptor–targeting domain, using established strategies for conditional cytokine activation and targeted delivery^38–40^.

Treatment prolonged survival in mice bearing WT BPPNM tumors, demonstrating that therapeutic engagement of the IFN-λ pathway can promote tumor control independently of *Bltp3a* loss (Fig. S4D).

We then examined whether BLTP3A dysfunction elicited comparable immune remodeling in human ovarian cancer systems. First, we determined that conditioned media from *BLTP3A*^KO^ OVCAR8 cells increased *CXCL9* and CD86 expression in THP-1-derived macrophages while limiting acquisition of a CD163^+^ CD206^+^ phenotype (Fig. 4E–G), consistent with the IFN-responsive macrophage polarization observed in our murine models. Recombinant IFN-λ similarly reduced CD163 expression (Fig. S4E), supporting direct macrophage responsiveness to this cytokine. To extend these observations *in vivo* we generated cord-blood-humanized NSG-SGM3 mice bearing HLA-A2-matched OVCAR3 tumors (Fig. 4H)^41,42^. Despite comparable total human CD45⁺ tumor infiltration, *BLTP3A* loss reduced the frequency of human CD68⁺CD163⁺ macrophages (Figs. S4F–H). Similarly, *BLTP3A^KO^* OVCAR3 tumors ectopically expressing M1098T exhibited fewer CD163⁺ macrophages and increased CD86⁺CD163⁻ macrophages relative to tumors overexpressing ancestral *BLTP3A* (*BLTP3A*^OE^) (Figs. 4I). Together, these findings support a model in which BLTP3A dysfunction promotes tumor-derived IFN-λ and chemokine production, fostering inflammatory macrophage remodeling and local CD8⁺ T-cell immunity (Fig. 4J).

### Ovarian cancer ascites induces a BLTP3A-regulated lysosomal stress–STING feedback circuit

Having established that BLTP3A dysfunction promotes IFN-λ-dependent antitumor immunity, we next sought to identify the cellular mechanism underlying this inflammatory response. Given BLTP3A’s role in endolysosomal remodeling and the chronic exposure of HGSOC cells to ascites^23,43^, we investigated whether the ovarian cancer microenvironment engages a BLTP3A-regulated lysosomal stress-response pathway. WT and *Bltp3a*^KO^ BPPNM cells were exposed to cell-free ascites and profiled by bulk RNA sequencing. In WT cells, ascites enriched pathways associated with proteasomal catabolism, endoplasmic- reticulum protein processing, MAPK and TNF signaling, and cellular stress responses (Fig. S5A), consistent with engagement stress-adaptation programs^16^. In contrast, under ascites exposure, *Bltp3a*^KO^ cells exhibited enrichment of glutathione metabolism, NRF2-mediated oxidative-stress responses, lipid- peroxidation responses, lysosomal pathways, and chaperone-mediated autophagy (Fig. 5A). Differential- expression analysis further revealed increased expression of lysosomal genes, including *Psap, Lamp1, Lamp2, Ctsb, Ctsl, Scarb2, Man2b1, Npc2, Ppt1, Hexa, and Hexb* (Fig. 5B) consistent with a coordinated lysosomal biogenesis and clearance program^44,45^. LysoTracker analysis revealed increased lysosomal staining in *Bltp3a*^KO^ BPPNM cells relative to WT cells under basal conditions and following treatment with LLOMe, a pharmacologic inducer of lysosomal membrane damage^46^, and ascites, consistent with expansion of the acidic lysosomal compartment (Fig. 5C). Both LLOMe and ovarian cancer ascites reduced inhibitory TFEB phosphorylation at Ser^122^, with a markedly greater reduction in *Bltp3a*^KO^ cells (Fig. 5D), indicating enhanced activation of lysosomal stress-response programs following BLTP3A loss^45,47–49^. These findings suggested an exaggerated lysosomal stress-adaptation response following BLTP3A loss. To determine whether this heightened lysosomal stress response was associated with increased membrane damage, we quantified galectin-3 (Gal3) puncta following LLOMe treatment. *Bltp3a*^KO^ cells exhibited greater Gal3 puncta accumulation than WT cells (Fig. S5B), consistent with increased lysosomal membrane damage. Notably, OVCAR3 cells expressing *BLTP3A*^M1098T^ similarly exhibited increased Gal3 puncta relative to cells expressing ancestral BLTP3A (Fig. 5E). Together, these findings suggest that ovarian cancer ascites induces lysosomal stress and that BLTP3A protects against lysosomal damage, a function that is compromised by the human M1098T variant.

**Figure 5.**
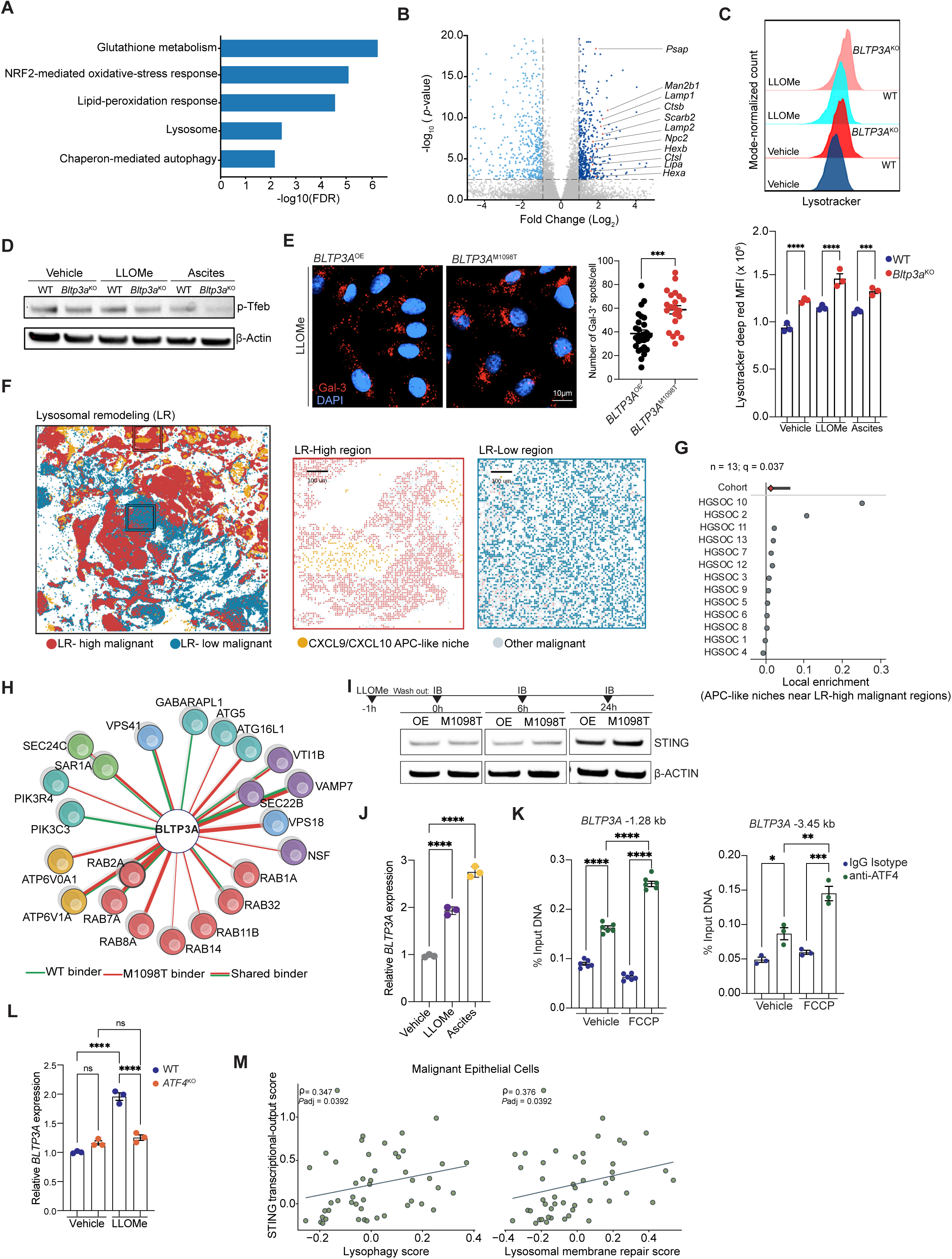
BLTP3A promotes resolution of lysosomal stress–induced STING signaling. **(A)** Gene-set enrichment pathways in ascites-exposed *Bltp3a*^KO^ versus WT BPPNM cells. **(B)** Volcano plot of differential gene expression in ascites-exposed *Bltp3a*^KO^ and WT BPPNM cells. Genes assigned to the KEGG lysosome gene set are highlighted, and selected lysosomal genes are labeled. **(C)** Representative LysoTracker fluorescence distributions and quantification of lysosomal signal in WT and *Bltp3a*^KO^ BPPNM cells treated as indicated. **(D)** Immunoblot of phospho-TFEB^Ser1^^22^ in WT and *Bltp3a*^KO^ cells treated with vehicle, LLOMe, or ascites. **(E)** Representative confocal images and quantification of Gal3^+^ puncta in *BLTP3A*^OE^ and *BLTP3A*^M1098T^ cells treated with LLOMe (1 mM, 1hr) and analyzed immediately after washout. **(F)** Visium HD epithelial regions classified as lysosomal remodeling-high or -low, with CXCL9/CXCL10-associated APC-like myeloid regions. (**G**) Local enrichment of CXCL9/CXCL10- associated myeloid regions near LR-high malignant regions across 13 HGSOC specimens (Cohort level q = 0.037). **(H)** BLTP3A immunoprecipitation–mass spectrometry interaction network; edge width indicates log₂ (IP/mock) enrichment and color indicates genotype specificity. **(I)** STING abundance following LLOMe treatment in *BLTP3A*^OE^ and *BLTP3A*^M1098T^ OVCAR3 cells. **(J)** Relative *BLTP3A* transcript expression in OVCAR8 cells treated with vehicle, LLOMe, or ovarian cancer ascites. **(K)** ATF4 chromatin immunoprecipitation-qPCR at candidate *BLTP3A* promoter sites located approximately 1.28 and 3.45 kb upstream of the transcriptional start site in vehicle or FCCP-treated OVCAR3 cells. **(L)** LLOMe-induced *BLTP3A* expression in WT and ATF4-deficient cells. **(M)** Spearman correlations between malignant cell STING-associated inflammatory transcriptional scores and lysophagy (ρ = 0.347) or lysosomal membrane-repair programs (ρ = 0.376) in HGSOC patient-level single-cell RNA-seq pseudobulks. Data are mean ± SEM unless otherwise indicated; each point represents an independent tumor or biological replicate. \**p* < 0.05, \*\**p* < 0.01, \*\*\**p* < 0.001; ns, not significant.

We next investigated whether lysosomal homeostasis and damage-response programs in malignant epithelial cells were associated with local immune organization in human HGSOC, using Visium HD spatial transcriptomic data from 13 tumor specimens. Tumor cell-enriched regions were classified using a composite lysosomal-remodeling transcriptional program encompassing lysosomal biogenesis, degradation, damage/repair, membrane protection, and lysophagy. Spatial mapping revealed heterogeneous expression of these programs within tumors, with representative high-scoring tumor regions exhibiting greater local enrichment of *CXCL9*^+^ inflammatory myeloid populations than low- scoring regions, an association observed across the cohort (Fig. 5F,G). These inflammatory myeloid regions were also associated with increased neighboring T cell abundance (Fig. S5C), suggesting a spatial relationship between epithelial lysosomal homeostasis, damage-response programs, and local myeloid–T cell organization.

We next sought to identify the molecular machinery through which BLTP3A regulates lysosomal stress adaptation. BLTP3A was recently identified as an effector of conjugation of ATG8 to single membranes (CASM), a noncanonical autophagy pathway involved in endolysosomal remodeling^23^. LC-MS/MS profiling of the BLTP3A interactome in OVCAR3 cells identified associations with RAB-family GTPases, V-ATPase subunits, PI3K complexes, HOPS/SNARE machinery, and ATG8/CASM-associated factors (Figs. 5H, S5D). Given the enrichment of oxidative-stress and lysosomal quality-control programs following ascites exposure (Fig. 5A-B), we next asked whether organelle stress influenced BLTP3A’s association with this trafficking network. Treatment with the mitochondrial uncoupler FCCP increased BLTP3A colocalization with RAB7-positive endolysosomal compartments (Fig. S5E), consistent with stress-dependent engagement of BLTP3A with late endolysosomal machinery. Interestingly, comparison of ancestral BLTP3A and M1098T interactomes revealed altered association with GABARAPL1 (Fig. 5H, S5D). This finding suggested that the M1098T variant may perturb the ability of BLTP3A to engage with ATG8-associated machinery involved in lysosomal stress responses.

Because STING engages CASM and facilitates TFEB-dependent lysosomal recovery, while its attenuation involves endolysosomal trafficking and processing^50–53^, we asked whether BLTP3A dysfunction impaired STING attenuation and thereby contributed to the inflammatory immune remodeling observed *in vivo*.

Following ascites exposure, STING protein remained elevated over 24 hours in *Bltp3a*^KO^ BPPNM cells (Fig. S5F). Direct STING activation with diABZI similarly prolonged STING accumulation in *Bltp3a*^KO^ BPPNM cells (Fig. S5G). Importantly, LLOMe-induced lysosomal injury similarly promoted sustained STING accumulation in OVCAR3 cells expressing *BLTP3A*^M1098T^ relative to ancestral BLTP3A (Fig. 5I). Together, these findings establish BLTP3A as a regulator of STING persistence following both direct pathway activation and lysosomal stress.

We next asked whether *BLTP3A* expression itself was regulated by lysosomal stress. We found that both ovarian cancer ascites and LLOMe increased *BLTP3A* transcript abundance (Fig. 5J). Given previous evidence that STING regulates ATF4-associated stress responses^54^ and that lysosomal perturbations can induce ATF4 independently of canonical integrated stress-response (ISR) signaling^55^, we investigated ATF4-dependent transcriptional regulation of *BLTP3A*. ChIP-qPCR demonstrated ATF4 occupancy at putative *BLTP3A* regulatory regions upstream of the transcriptional start site, with increased enrichment following FCCP treatment (Fig. 5K). Supporting this, genetic ablation of ATF4 markedly impaired LLOMe-induced *BLTP3A* expression (Fig. 5L). In contrast, ISRIB treatment did not prevent this induction (Fig. S5H), supporting prior findings that lysosomal injury engages an ATF4-dependent transcriptional response that is insensitive to canonical ISR inhibition^55^. Interestingly, we next found that deletion of *Sting1* attenuated LLOMe and Ascites-induced *Bltp3a* expression in BPPNM cells (Fig. S5I). Together, these findings support a negative-feedback circuit in which lysosomal stress promotes STING-dependent, ATF4-mediated *BLTP3A* induction, while BLTP3A in turn limits STING persistence following stress- induced activation.

Finally, we examined whether lysosomal remodeling was associated with STING-related inflammatory signaling in human HGSOC. In malignant HGSOC cells analyzed within an integrated ovarian cancer single-cell RNA-sequencing resource^28^, we determined that STING-associated inflammatory signatures correlated with lysophagy (ρ = 0.347) and lysosomal membrane-repair programs (ρ = 0.376) (Fig. 5M). Together with our spatial transcriptomic findings, these observations support a relationship between lysosomal remodeling, STING-associated inflammatory signaling, and local immune organization in human HGSOC, consistent with the STING–BLTP3A feedback circuit identified in our experimental models.

### BLTP3A governs LC3/CASM-associated lysosomal stress resolution to attenuate STING signaling

Having established that BLTP3A limits STING protein persistence, we next investigated whether its loss also prolonged STING signaling following lysosomal injury. LLOMe induced phosphorylation of STING^Ser3^^65^ and TBK1^Ser1^^72^ in WT cells, whereas *Bltp3a*^KO^ cells exhibited greater and more persistent phosphorylation of both proteins during recovery (Fig. 6A), demonstrating impaired attenuation of STING signaling.

**Figure 6.**
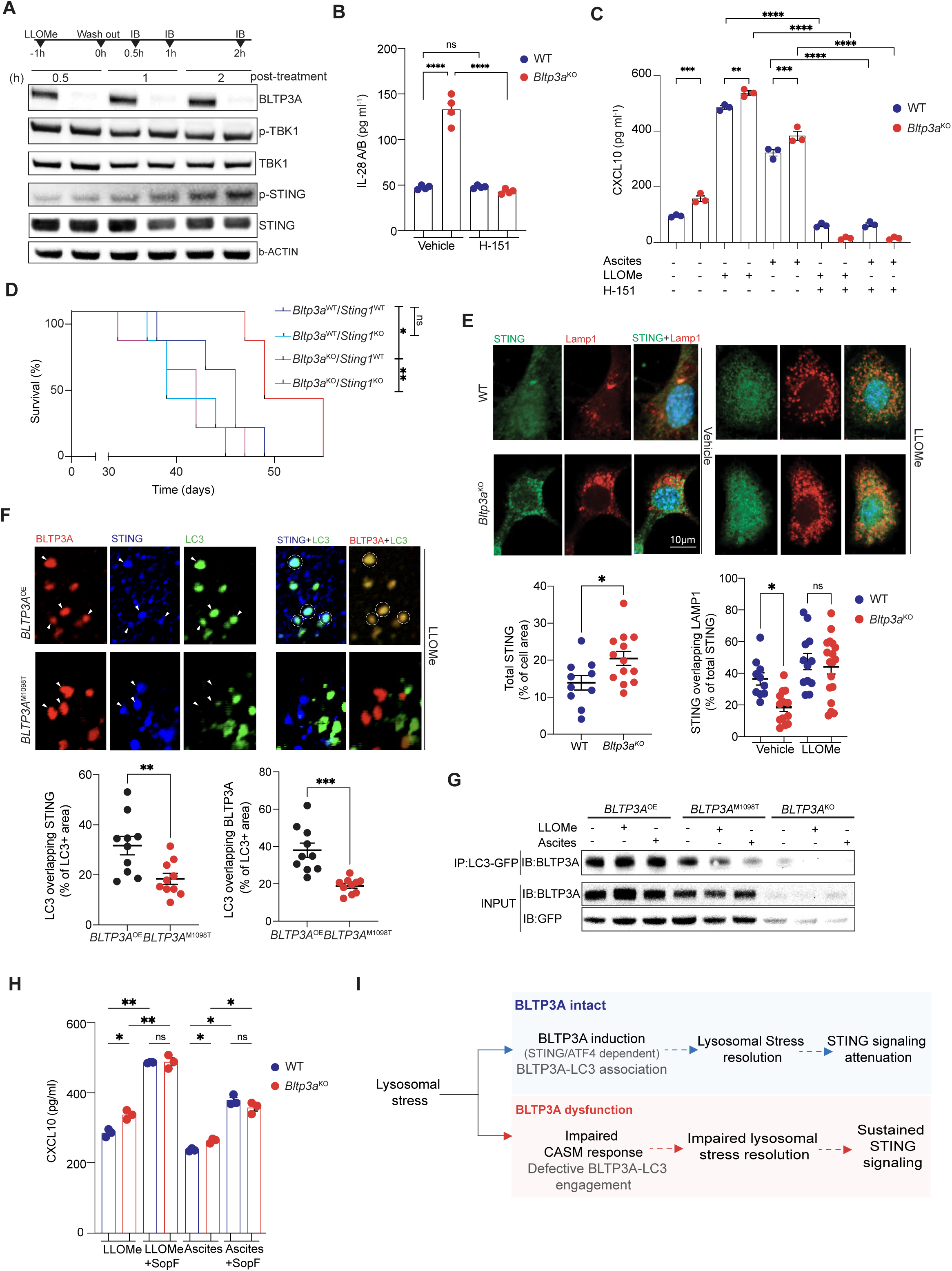
BLTP3A resolves lysosomal STING signaling through ATG8/CASM-associated remodeling. **(A)** pSTING^Ser365^ and pTBK1^Ser172^ phosphorylation and total protein abundance following an hour LLOMe pulse and washout in WT and *Bltp3a*^KO^ cells. **(B)** IFN-λ2/3 secretion with or without STING antagonist H-151. **(C)** CXCL10 secretion following vehicle, ascites, or LLOMe treatment, with or without H-151. **(D)** Survival of mice bearing BPPNM tumors of the indicated *Bltp3a* and *Sting1* genotypes (n= 5 mice per group). **(E)** Representative confocal images of STING and LAMP1 in WT and *Bltp3a*^KO^ BPPNM cells treated with vehicle or LLOMe. Quantification shows the total STING-positive area per cell under vehicle condition (left) and the fraction of total STING overlapping Lamp1 in vehicle- and LLOMe-treated cells (right). **(F)** Representative confocal images of LLOMe-treated *BLTP3A*^OE^ and *BLTP3A*^M1098T^ OVCAR8 cells stained for LC3, BLTP3A and STING represented by Arrowheads, and regions of their colocalization indicated by dashed circles. Quantification shows the percentage of LC3-STING (left) and LC3-BLTP3A (right) overlapping area **(G)** LC3-GFP immunoprecipitation and input immunoblots from *BLTP3A*^OE^ and *BLTP3A*^M1098T^ OVCAR8 cells following vehicle, LLOMe, or ascites exposure. **(H)** CXCL10 concentrations in conditioned media from WT and *Bltp3a*^KO^ BPPNM cells exposed to LLOMe or ascites, with or without SopF ectopic expression. **(I)** Proposed BLTP3A–ATG8/CASM–STING negative-feedback model. Data are mean ± SEM unless otherwise indicated; each point represents an independent tumor or biological replicate. Survival curves were compared by log-rank tests; other comparisons used the tests described in Methods. \**p* < 0.05, \*\**p* < 0.01, \*\*\**p* < 0.001; ns, not significant.

We next asked whether this sustained STING activity was responsible for the inflammatory secretory program and antitumor immunity elicited by BLTP3A deficiency. Pharmacologic STING antagonism with H-151 abolished the elevated secretion of IFN-λ2/3 (IL-28A/B) and markedly suppressed LLOMe- and ascites-induced CXCL10 production in *Bltp3a*^KO^ cells (Figs. 6B, C). Intracellular cGAMP was not increased in *Bltp3a*^KO^ cells relative to WT BPPNM cells following either treatment (Fig. S6A), consistent with regulation of STING signaling persistence rather than increased upstream agonist production. Importantly, deletion of *Sting1* in *Bltp3a*^KO^ BPPNM tumor cells abolished the survival advantage in syngeneic systems (Fig. 6D), establishing a requirement for tumor cell-intrinsic STING in the protective antitumor response.

Because STING signaling is terminated through post-Golgi endolysosomal trafficking and processing^56,57^, we examined whether BLTP3A influenced the intracellular fate of STING. Under basal conditions, *Bltp3a*^KO^ cells exhibited increased STING puncta relative to WT cells, with limited overlap between the excess STING puncta and LAMP1-positive compartments (Fig. 6E). Following LLOMe treatment, however, STING redistributed to LAMP1-positive compartments to a similar extent in WT and *Bltp3a*^KO^ cells, despite its persistent accumulation in *Bltp3a*^KO^ cells. LLOMe also increased STING accumulation within TGN46-positive compartments in *Bltp3a*^KO^ cells (Fig. S6B). Together with the prolonged STING accumulation following LLOMe-induced lysosomal injury in *BLTP3A*^M1098T^ expressing cells (Fig. 5I), these findings suggest that BLTP3A regulates STING persistence following activation without preventing its redistribution to post-Golgi and endolysosomal compartments.

Given that STING activation promotes V-ATPase–ATG16L1-dependent lipidation of ATG8-family proteins, including LC3, on single-membrane compartments^51^, we investigated whether the M1098T variant disrupted the stress-dependent association of BLTP3A and STING with LC3-positive membranes. Following LLOMe exposure, cells expressing *BLTP3A*^M1098T^ displayed reduced LC3–STING and BLTP3A–LC3 colocalization relative to cells overexpressing ancestral BLTP3A (*BLTP3A*^OE^) (Fig. 6F). These findings suggest that M1098T variant disrupts the engagement of BLTP3A and STING-containing compartments with LC3-positive membranes during lysosomal stress. We therefore directly examined the stress–regulated BLTP3A–LC3 association. Under basal conditions, both BLTP3A^OE^ and BLTP3A^M1098T^ co-immunoprecipitated with LC3 (Fig. 6G). Following LLOMe or ascites exposure, however, LC3 association increased with ancestral BLTP3A but was markedly reduced with BLTP3A^M1098T^. Thus, BLTP3A^M1098T^ preserves basal LC3 binding but selectively disrupts stress-enhanced BLTP3A–LC3 engagement. Together, these findings demonstrate that the M1098T variant selectively impairs stress- enhanced BLTP3A–LC3 engagement, accompanied by reduced STING–LC3 colocalization, suggesting that BLTP3A participates in the association of STING-containing compartments with ATG8-associated endolysosomal remodeling machinery.

To determine whether CASM-associated machinery contributed to BLTP3A-dependent inflammatory regulation, we expressed SopF in BPPNM cells to disrupt V-ATPase–ATG16L1 coupling^51^. Following LLOMe or ascites exposure, SopF increased CXCL10 production and eliminated the difference between WT and *Bltp3a*^KO^ cells (Fig. 6H). Together with the impaired stress-dependent LC3 engagement observed with BLTP3A^M1098T^, these findings implicate a SopF-sensitive, CASM-associated process in STING attenuation (Fig. 6I). Thus, BLTP3A functions as a stress-inducible regulator of STING persistence and inflammatory output, providing a mechanistic basis for the protective myeloid–CD8⁺ T cell immunity elicited by BLTP3A dysfunction.

## DISCUSSION

Here, we identify BLTP3A as a tumor cell-intrinsic lysosomal stress-response regulator that limits the duration and inflammatory output of STING signaling in ovarian cancer. The germline *BLTP3A*^M1098T^ variant was associated with improved HGSOC survival and spatial organization of cytotoxic T cells with inflammatory macrophages. *Bltp3a* loss in tumors cells promoted STING-dependent IFN-λ and chemokine production, inflammatory myeloid remodeling, and protective CD8⁺ T cell immunity. Notably, M1098T reproduced key features of the macrophage and lysosomal stress phenotypes observed following BLTP3A deficiency, connecting the human genetic association to the cellular mechanism uncovered in our experimental models.

STING signaling is closely integrated with lysosomal homeostasis: activated STING promotes TFEB- dependent lysosomal biogenesis and V-ATPase–ATG16L1-dependent CASM, while its attenuation involves endolysosomal trafficking and processing^51,53,58,59^. Lysosomal membrane injury can also activate cGAS–STING-dependent type I interferon signaling through mitochondrial DNA release^60^. Recent studies have further shown that disruption of STING trafficking or degradation can prolong signaling and enhance antitumor immunity^59,61^. Our findings identify BLTP3A as an endogenous regulator of this stress-response pathway. Lysosomal stress induced *BLTP3A* through STING-dependent, ATF4-mediated transcription, while BLTP3A deficiency prolonged STING–TBK1 signaling despite preserved STING redistribution to LAMP1-positive compartments. Moreover, lysosomal stress enhanced BLTP3A association with LC3, whereas the M1098T variant compromised this response. Together with the BLTP3A interactome and functional disruption of CASM-associated machinery, these findings support a model in which BLTP3A participates in ATG8/CASM-associated lysosomal remodeling to constrain STING signaling. Thus, impaired BLTP3A-dependent stress resolution sustains tumor-cell inflammatory output and promotes the protective myeloid–T cell immunity identified in our experimental models.

IFN-responsive myeloid cells can form intratumoral niches that recruit, activate, and retain antitumor T cells, whereas tumor-intrinsic MAPK hyperactivation and PGE_2_ programs can disrupt this organization^8,10^. Less is known about tumor-autonomous pathways that actively establish such niches. Our data identify failed lysosomal stress resolution as one such mechanism: persistent tumor-cell STING signaling sustained IFN-λ and chemokine production, remodeled the myeloid compartment toward IFN-responsive macrophage states, and increased their spatial association with effector T cells. Importantly, these myeloid niches supported local CD8^+^ T cell proliferation and maintenance, providing a functional basis for the cytotoxic T cell–myeloid organization associated with the protective BLTP3A M1098T variant. This immune program depended on tumor-derived IFN-λ rather than type I interferon. This bias may reflect the epithelial lineage of these tumors, as type III interferons are preferentially secreted at epithelial interfaces and can be induced downstream of STING signaling^62,63^. Tumor-derived IFN-λ3 was recently shown to enhance macrophage phagocytosis, proinflammatory polarization, cytotoxic lymphocyte recruitment, and responsiveness to immune-checkpoint blockade in bladder cancer^64^. Our findings extend this paradigm by identifying lysosomal stress resolution as an upstream determinant of tumor-derived IFN-λ and showing how a tumor-cell organelle-stress program can establish myeloid niches that locally sustain antitumor T cell immunity.

Advanced HGSOC develops within a lipid-rich peritoneal ascites environment that imposes chronic tumor cell stress. We found that ascites expanded and remodeled the lysosomal compartment, altered TFEB signaling, and induced BLTP3A. These findings identify ovarian cancer ascites as a driver of tumor cell lysosomal stress and place lysosomal stress resolution within the adaptive response to the peritoneal microenvironment. Recent studies have similarly shown that ovarian cancer ascites reprograms tumor cell lipid and redox homeostasis, promoting lipid storage and resistance to ferroptosis^17^. Our findings add lysosomal integrity and stress recovery to this adaptive program and connect ascites-induced lysosomal remodeling to tumor-intrinsic innate immune signaling. Consistent with this model, lysosome-related ovarian cancer states have been associated with distinct interferon programs, macrophage composition, CD8^+^ T cell infiltration, and clinical outcome^65^. Our findings provide a mechanistic framework for this association by connecting tumor cell lysosomal stress resolution to STING persistence and downstream myeloid–T cell immunity. Consistent with ongoing lysosomal stress adaptation in human disease, primary HGSOC malignant epithelial cells exhibited coordinated lysosomal remodeling and STING-associated transcriptional programs, suggesting that this pathway operates within the native tumor microenvironment.

Our findings suggest that the downstream IFN-λ axis may be therapeutically actionable. The antitumor activity of our tumor-targeted IFN-λ procytokine provides proof of principle for therapeutic engagement of this pathway, with conditional cytokine activation offering a potential strategy to limit systemic exposure. More broadly, defining the molecular steps governing BLTP3A-dependent lysosomal stress recovery may reveal additional strategies for sustaining endogenous, stress-induced STING activity and its antitumor inflammatory output.

Several mechanistic questions remain. The precise BLTP3A-dependent step in lysosomal stress resolution remains undefined. Recent studies have established ER–lysosome membrane contacts and lipid transfer as important mechanisms of lysosomal repair^66^, and a similar function has been proposed for BLTP3A following its CASM-dependent recruitment to damaged lysosomes^23^. One possibility is that stress- enhanced LC3 engagement positions BLTP3A to facilitate membrane lipid transfer and lysosomal recovery. Whether the M1098T variant disrupts this process, and how such a defect might contribute to STING persistence, remain important questions.

## EXPERIMENTAL MODELS

### Animal experiments and ethics

Mice: All animals were maintained and used in accordance with the Institutional Care and Use Guidelines of Rutgers University (IACUC protocols # 202100080). All mice are housed in pathogen-free conditions with a housing temperature of 22 ± 1°C, 55 ± 5% humidity and a photoperiod of 14 h of light and 10 h of dark. Within each experiment, age-matched female mice were used. C57BL/6, OT-1 mice were purchased from Charls River, then bred in-house. *NOD.Cg Prkdc^scid^ Il2rg^tm1Wjl^ Tg(CMV-IL3, CSF2, KITLG)1Eav/MloySzJ* (NSG-SGM3), *NOD.Cg-Prkdc^scid^ Il2rg^tm1Wjl^/SzJ* (NSG), *B6.129S(C)-Batf3^tm1Kmm^/J* (*Batf3*^-/-^) mice, and *B6.Cg-Rag2^tm1.1Cgn^/J* (*Rag2*^-/-^) mice were purchased from the Jackson Laboratory, then bred in-house. For *Rag2*^-/-^*Batf3*^-/-^ strain generation, *Batf3*^-/-^ mice were crossed to *Rag2*^-/-^ mice, and homozygous offspring (*Rag2*^-/-^ x *Batf3*^-/-^) were confirmed by genotyping. For all the above strains, mice were 7-12 weeks old. For OT-1 CD8^+^ T cell isolation, 8- 20-week-old OT-1 mice were used. Mice were euthanized when the human end point was reached, when tumors reached the maximum allowed tumor volume of 1500 mm^3^. Autochthonous ovarian tumors were induced in *p53^f/f^* ^LSL^*Kras*^G12D^ and *p53^f/f^*^LSL^*Kras*^G12D^ *Bltp3a^f/f^* transgenic mice through delivery of adenoviral-expressing Cre recombinase (U of Iowa) into the ovarian bursa, as previously described^67^.

### Human Specimens

Fresh human ovarian carcinoma tissues, ascites fluid, and peripheral whole blood were procured under protocols approved by the Institutional Review Board (IRB) at Rutgers University Office for Research (Study ID: Pro2022000213); and under a protocol approved by the Biospecimen Repository Service (BRS) at the Rutgers Cancer Institute (#002201). Archived specimens including histopathology slides, tumor tissue microarrays, and Formalin-Fixed Paraffin-Embedded (FFPE) scrolls were acquired under the same BRS protocol (#002201). Informed consent was obtained from all subjects. No protected health information was received.

Human peripheral whole blood bags were procured from the New York Blood Center (Long Island City, NY) under a protocol approved by the IRB at Rutgers University Office for Research (Study ID: Pro2021002369). Human cord blood was also procured from the New York Blood Center under a separate protocol approved by the IRB at Rutgers University Office for Research (Study ID: Pro2022000577).

Fresh tumor chunks were flash frozen for downstream protein, DNA, and RNA isolation. Cord blood was processed using a Ficoll density gradient protocol to obtain PBMCs and CD34^+^ cells were isolated using a human CD34 MicroBead Kit (Miltenyi; 130-046-702).

### Genetic Tumor Models and Cell Lines

ID8 cells were retrovirally transduced to express *Defb29* and *Vegfa*, as previously described^68^. ID8^Defb29/Vegfa^ cells were implanted intraperitonially into syngeneic C57/B6 mice to generate ascites fluid, which was collected after 45-50 days and centrifuged to produce cell-free ascites. Aliquots were stored at -80°C and subsequently used in downstream experiments as described. HEK 293T/17 cells, OVCAR8, OVCAR3 were purchased from ATCC (Manassas, VA) and cultured in DMEM (Gibco). THP-1 cell line was purchased from ATCC and cultured in RPMI-1640 (Gibco). BPPNM cells were a gift from Dr. Robert Weinberg^29^ and were maintained in DMEM with 4% FBS, 1% ITS, 2 ng/ml epidermal growth factor (EGF) (Sigma E9644-.2MG) and 1% penicillin-streptomycin. All media for cell lines were supplemented with 10% FBS, 2 mM L-glutamine (Gibco) and 100 IU ml^-1^ penicillin-streptomycin, 1x sodium pyruvate (Thermo Fischer). CD8^+^ T cells were cultured in full T cell medium containing RPMI-1640 supplemented with 10% FBS, 2 mM L-glutamine (Gibco) and 100 IU ml^-1^ penicillin-streptomycin, 1x sodium pyruvate and 0.05 mM β-mercaptoethanol (Millipore). All cells were cultured at 37C and 5% CO2. Cells routinely tested negative for mycoplasma contamination.

### Genetic Engineering of Tumor Cell Lines

BLTP3A, STING, ATF4, and Ifnl2/3-deficient tumor cell lines were generated by CRISPR–Cas9 ribonucleoprotein (RNP) electroporation using the Neon Transfection System (Invitrogen; NEON1S). OVCAR3 and OVCAR8 cells were targeted for *BLTP3A*, or *ATF4*, and BPPNM cells were targeted for *Bltp3a*, *Sting1*, or *Ifnl2/3*, as indicated.

For crRNA-based targeting, custom crRNAs (IDT) were duplexed with tracrRNA (IDT; 1072534) at equimolar concentrations by heating to 95°C for 5 min followed by gradual cooling to room temperature. The resulting crRNA duplexes were combined with Alt-R S.p. Cas9 Nuclease V3 (IDT; 1081059) at equimolar concentrations and incubated for 10 min at room temperature to generate RNP complexes. For sgRNA-based targeting, custom sgRNAs (GenScript) were combined with Alt-R S.p. Cas9 Nuclease V3 at a 3:1 molar ratio (sgRNA) and incubated for 20 min at room temperature. RNP complexes were delivered by electroporation using the Neon Transfection System at 1,170 V, 30 ms, and 2 pulses. Cells were subsequently cultured in complete medium and expanded. Where necessary, electroporation was repeated for two to three rounds to achieve efficient target ablation. Loss of *BLTP3A*, *ATF4, Bltp3a*, *Sting1* was confirmed by immunoblotting, whereas loss of IFN-λ 2/3 production in *Ifnl2/3* -targeted cells was validated by ELISA. crRNA sequences are provided below.

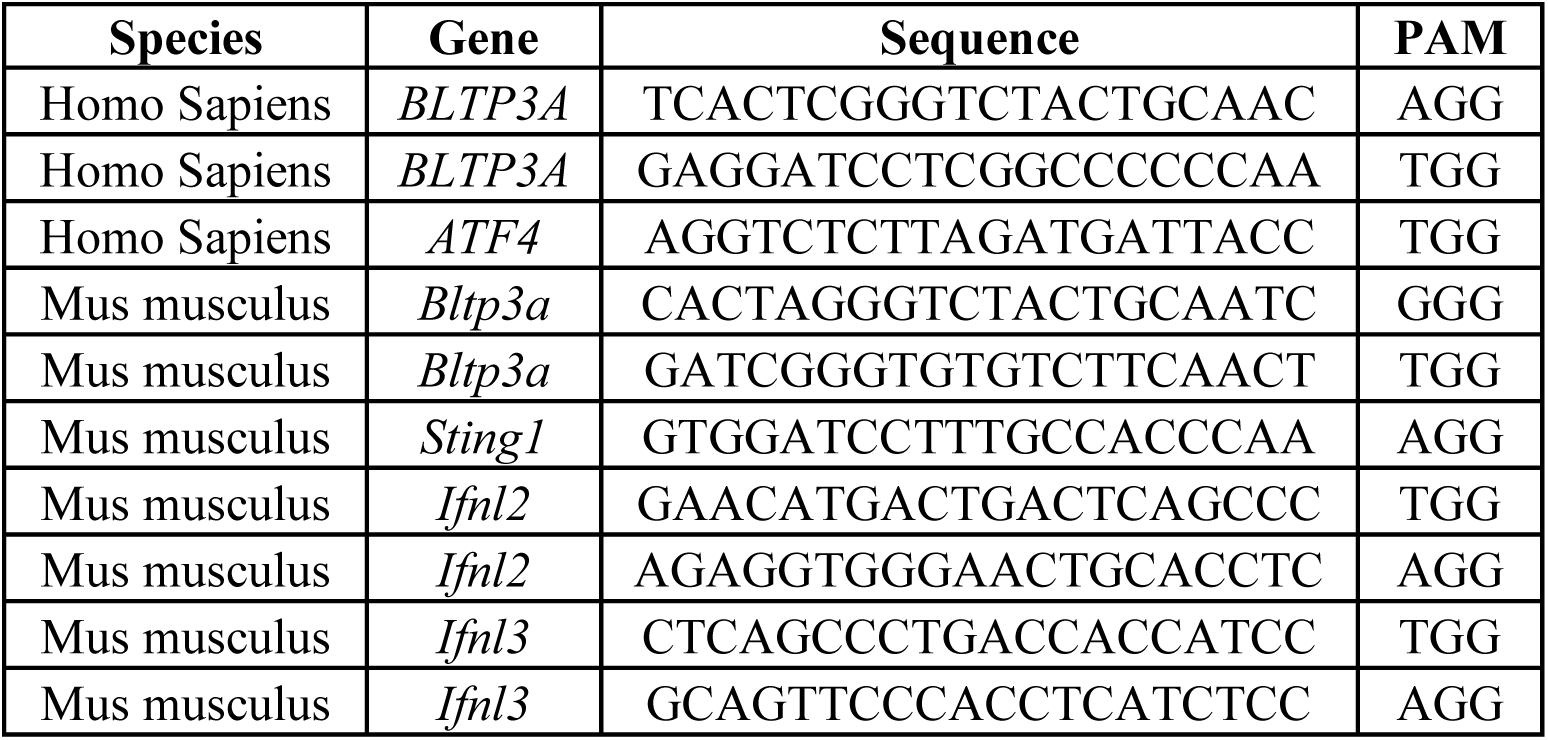

For reconstitution experiments, BLTP3A-deficient OVCAR3 and OVCAR8 cells were transduced with pLVX-IRES-ZsGreen (Takara Bio) encoding C-terminal FLAG-tagged ancestral *BLTP3A*, *BLTP3A*^M1098T^, or empty vector. Lentiviral particles were generated in Lenti-X 293T cells by cotransfection of the corresponding transfer vector with psPAX2 and pMD2.G using jetPRIME. Virus- containing supernatants were collected 48–72 h after transfection, clarified, passed through a 0.45 μm filter, and used directly or concentrated using Lenti-X Concentrator. Target cells were transduced in the presence of polybrene, with centrifugation-enhanced transduction where indicated. Transduction efficiency was assessed by ZsGreen expression using flow cytometry.

BPPNM cells were additionally transfected with the SOPF plasmid (Addgene: #135174) using jetPRIME according to the manufacturer’s instructions and subsequently cultured in complete medium before use in downstream experiments.

Plasmids were propagated in NEB 5-alpha competent *Escherichia coli* and purified using Qiagen plasmid purification kits. pcDNA3.1-BLTP3A-FLAG was generated by EcoRI/NotI subcloning from the corresponding pLVX donor vector and verified by restriction analysis.

### *In vivo* mouse studies

For the generation of orthotopic BPPNM tumors, 3x 10^6^ cells per 100ul PBS and were mixed 1:1 with 100ul Matrigel (Corning) for a total of 200ul per interperitoneal (i.p.) injection. Matrigel/cell aliquots were individually prepared for each tumor injection to decrease variability of tumor injection. Anti-CD8 (Clone 2.43), BioXCell) depleting antibodies were delivered i.p. at 150 µg per dose. Treatment commenced 3 days prior to tumor engraftment, was administered again on the day of engraftment, and was maintained at 3-day intervals thereafter until day 21. Anti-CXCR3 (CXCR3-173, BioXCell) monoclonal antibodies were given at 200µg/ dose and administered i.p., starting from day 5 and was maintained at 4-day intervals thereafter until day 24. Isotype controls were used to confirm the lack of non-specific effects. Anti-Gr-1 (RB6-8C5) monoclonal antibodies were given at 150µg/dose and administered intraperitoneally every two days. Anti-IFNAR1 blocking antibody (clone MAR1-5A3; Bio X Cell, cat. no. BE0241) was administered intraperitoneally at 200 µg per mouse per dose. Treatment began on the day of tumor challenge and continued at 4-day intervals until the experimental endpoint. To assess the proliferative activity of tumor- infiltrating OT-I T cells during inhibition of lymph-node egress, mice received FTY720 (Sigma) intraperitoneally at 20 µg per mouse in 100 µL saline. Treatment began on the day of OT-I T cell transfer and continued daily for four consecutive days. For OVCAR3 cell lines, 5x 10^6^ cells per 100ul PBS and were mixed 1:1 with 100ul Matrigel for a total of 200ul per interperitoneally injection in humanized NSG mice. Intraperitoneal Tumor measurements were recorded using BLI-IVIS imaging with prior luciferin injection. Mice bearing luciferase-expressing tumors received D-luciferin at 150 mg/kg intraperitoneally and were imaged after 10 minutes under isoflurane anesthesia. Signal was quantified as total photon flux using Living Image software. Mice were euthanized at protocol-defined humane endpoints based on tumor burden, ascites accumulation, impaired mobility or respiration, or other signs of morbidity.

### DNA-encoded tumor-targeted IFN λ procytokine treatment

A DNA-encoded, tumor-targeted IFN λ procytokine was designed to enable conditional delivery of murine IL-28 (IFN λ2/3) within the tumor microenvironment. The construct incorporated an FSH receptor (FSHR)-targeting domain to promote tumor localization, matrix metalloproteinase (MMP)-cleavable elements to enable protease-dependent activation within the tumor microenvironment, and an XTEN domain to extend circulating half-life. The complete procytokine sequence was synthesized by GenScript and cloned into the pVAX1 expression plasmid. Empty pVAX1 plasmid was used as the corresponding control.

For in vivo gene delivery, tumor-bearing mice received 80 μg of IFN λ procytokine-encoding or control pVAX1 plasmid in a total volume of 20 μL by intratibial injection, followed immediately by electroporation using the AgilePulse ID In Vivo Gene Delivery System (BTX). Electroporation was performed using a four-needle array with a 4 mm inter-row spacing and 3 mm needle length. For the 4 mm electrode configuration, pulses consisted of one 450 V pulse for 0.05 ms, a second 450 V pulse for 0.05 ms, and eight 110 V pulses for 10 ms, with pulse intervals of 0.2 ms, 50 ms, and 20 ms for the respective pulse groups. Mice received two treatments, with the first administered 14 days after tumor inoculation and the second administered 10 days later.

### Generation of humanized mice and human EOC xenografts

Humanized mice were generated by reconstituting sublethally irradiated NSG-SGM3 mice with human cord blood-derived HLA-A2^+^ CD34⁺ hematopoietic stem and progenitor cells (HSPCs). CD34⁺ HSPCs were isolated from human cord blood by magnetic bead-based enrichment using a CD34 MicroBead Kit UltraPure, human (# 130-100-453) according to the manufacturer’s instructions. Recipient mice received 250 cGy total-body irradiation followed by retro-orbital administration of 1 ×10⁵ purified CD34⁺ HSPCs. Mice were allowed to undergo human hematopoietic reconstitution for 8–12 weeks before tumor challenge. Where indicated, peripheral-blood engraftment was assessed before tumor implantation by flow cytometric quantification of human CD45⁺ leukocytes. Reconstituted mice were challenged intraperitoneally with 5 × 10⁶ of WT, *BLTP3A*^KO^, *BLTP3A*^OE^, *BLTP3A*^M1098T^ OVCAR3 human ovarian cancer cells, with three female mice per group and monitored for tumor progression. Endogenous β2m was CRISPR/Cas9 ablated from these OVCAR3 cells, followed by lentiviral rescue with a single-chain HLA-A2-β2m fusion to reduce allogenic activity^69^. Mice were euthanized 10 weeks after tumor-cell implantation, and tumors and associated tissues were collected for downstream flow cytometric and molecular analyses.

### *In vivo* ovalbumin antigen-presentation assay

To assess antigen presentation by tumor-associated macrophages, C57BL/6 mice bearing BPPNM WT or *Bltp3a*^KO^ tumors received 200 μg EndoGrade ovalbumin (OVA; LIONEX, cat. no. LET0027) in 100 μL PBS by intraperitoneal injection on day 34 after tumor challenge. Tumors were harvested 24 h after OVA administration and processed into single-cell suspensions. Tumor-associated macrophages were identified as CD11b⁺F4/80⁺MHC-II⁺ cells and subdivided into CD86⁺CD206⁻ and CD86⁻CD206⁺ populations. Surface SIINFEKL–H-2Kᵇ complexes were assessed by flow cytometry using a PE-conjugated anti- mouse H-2Kᵇ–SIINFEKL antibody (clone 25-D1.16; BioLegend).

### In Vivo Analysis of pMHCI Cross-Dressing on Macrophages

To establish an MHCI-mismatched haplotype model, 3 × 10^6^ *Bltp3a*^KO^ BPPNM cells (C57BL/6 origin; H-2K^b^) were injected intraperitoneally into BALB/c (H-2K^d^) hosts, as defined by others^14^. To preclude T cell-mediated allograft rejection, mice were administered by an anti-CD8 depleting antibody (clone 2.43, BioXCell) or a rat IgG2b isotype control. Antibodies were delivered *i.p.* at 150 µg per dose. Treatment commenced 3 days prior to tumor engraftment, was administered again on the day of engraftment, and was maintained at 3-day intervals thereafter until day 21. Thirty-five days post-inoculation, excised tumors were processed into single-cell suspensions for direct flow cytometric detection of H-2K^b^.

### Design and construction of an FSHR-directed IFN-λ3 procytokine expression vector

A chimeric type III interferon procytokine was designed to enable tumor-directed delivery of IFN-λ3 through follicle-stimulating hormone receptor (FSHR) targeting and matrix metalloproteinase (MMP)- dependent proteolytic processing. The fusion protein consisted of an N-terminal follicle-stimulating hormone β subunit (FSHβ), the common glycoprotein hormone α subunit (CGA), the mature murine IFN- λ3 (IL-28B) sequence (NCBI RefSeq NP_796370.1, residues 20–193), and a C-terminal XTEN144 polypeptide intended to prolong circulating half-life. The FSHβ and CGA domains were connected by a flexible (Gly₄Ser)₃ linker to support FSHR-directed targeting. A second (Gly₄Ser)₃ linker separated CGA from an MMP-sensitive cleavage sequence (GPLGVR), followed by a (Gly₄Ser)₂ linker and the IFN-λ3 domain. The cytokine domain was further connected to XTEN144 through a (Gly₄Ser)₂ linker, a second MMP-sensitive cleavage sequence (GPLGSR), and another (Gly₄Ser)₂ linker. The resulting construct had the following N- to C-terminal organization: FSHβ–(Gly₄Ser)₃–CGA–(Gly₄Ser)₃–GPLGVR–(Gly₄Ser)₂– IFN-λ3–(Gly₄Ser)₂–GPLGSR–(Gly₄Ser)₂–XTEN144.

The two MMP-sensitive linkers were incorporated to facilitate proteolytic processing of the cytokine- containing fusion protein within the tumor microenvironment, while the gonadotropin-derived targeting domains were intended to promote localization to FSHR-expressing ovarian tumor cells. The XTEN144 domain was included to increase the hydrodynamic size and prolong systemic persistence of the fusion protein. The chimeric coding sequence was incorporated into the pVAX1 mammalian expression vector (Thermo) to enable expression of the engineered procytokine following plasmid delivery.

## METHOD DETAILS

### Tumor dissociation and cell isolation

Murine tumors were measured, dissected free of surrounding adipose tissue and mesenteric lymph nodes, weighed, and processed into single-cell suspensions. Tumors were enzymatically and mechanically dissociated in RPMI-based tumor dissociation medium containing enzymes from the mouse tumor dissociation kit (the Miltenyi mouse tumor dissociation kit) using a gentleMACS tissue dissociator with heaters and the murine TDK-2 program. Dissociated tumors were filtered through a 100-μm cell strainer, washed, and resuspended in calcium- and magnesium-free PBS at a volume normalized to tumor weight. For flow cytometry staining, 100 μL of the resulting cell suspension was used per tumor sample. For bone marrow isolation, femurs and tibias were collected and flushed with 5 mL of complete RPMI medium supplemented with 10% FBS, 2 mM L-glutamine, 1 mM sodium pyruvate, and 100 U/mL penicillin- streptomycin. Bone marrow suspensions were filtered through a 70-μm cell strainer. Red blood cells were lysed with 1–2 mL RBC lysis buffer for 2–4 min at room temperature, and lysis was quenched with PBS containing 2% FBS. Cells were then washed, counted, and used for bone marrow chimera experiments. For splenocyte preparation, spleens were mechanically dissociated using a glass tissue grinder, followed by red blood cell lysis in 5–10 mL RBC lysis buffer for 2–4 min at room temperature. Lysis was quenched with PBS containing 2% FBS, and cells were washed, filtered as needed, counted, and used for downstream flow cytometry, sorting, or functional assays. All centrifugation steps were performed at 500 × g for 4–5 min at 4°C. Naïve CD8⁺ T cells were isolated from OT-I mouse (JAX: #003831) spleens by magnetic enrichment using the MagniSort Mouse Naïve CD8⁺ T Cell Enrichment Kit according to the manufacturer’s instructions.

### Cell treatments

Cells were treated with the STING agonist diABZI (1 µM; MedChemExpress) or the STING antagonist H-151 (2 µM; MedChemExpress) overnight, unless otherwise indicated. To induce lysosomal membrane damage, BPPNM, OVCAR8, and OVCAR3 cells were treated with L-leucyl-L-leucine methyl ester (LLOMe; MedChemExpress) at concentrations of 1 mM and 3 mM, and 1 mM, respectively. For cytokine analysis, BPPNM cells were exposed to LLOMe for 1 h, washed, and incubated overnight in fresh culture medium before supernatant collection. For immunoblotting and confocal microscopy, treatment durations and recovery periods varied according to the experimental conditions specified in the corresponding Results sections.

### Antibodies and flow cytometry

Tumors were harvested 35 days after tumor implantation and processed into single-cell suspensions as described above. Cells were first stained with Zombie NIR Fixable Viability Dye (BioLegend) for 10 min at room temperature to enable exclusion of dead cells. Following washing, cells were incubated with purified anti-mouse CD16/32 Fc block (BioLegend; 1:100) to minimize nonspecific Fc receptor-mediated antibody binding. Surface antibodies were diluted in FACS buffer consisting of PBS supplemented with 2% FBS and 2 mM EDTA, and cells were stained for 20–25 min at 4°C in the dark. Cells were subsequently washed in FACS buffer by centrifugation at 500 × g for 5 min at 4°C. For intracellular staining, cells were fixed and permeabilized using the Foxp3/Transcription Factor Staining Buffer Set (eBioscience/Thermo Fisher Scientific) according to the manufacturer’s instructions. Intracellular antibodies were diluted in permeabilization buffer and incubated with cells according to the manufacturer- recommended staining conditions. Fluorescence-minus-one (FMO) controls, single-color compensation controls, and unstained controls were included as appropriate to establish gating boundaries and fluorescence compensation. For analysis, debris, doublets, and nonviable cells were excluded before identification of the indicated immune-cell populations. Flow cytometry data were analyzed using FlowJo software (BD Biosciences).

### Murine EOC multiplexed immunofluorescence and image analysis

Formalin-fixed, paraffin-embedded (FFPE) mouse tumor tissue microarrays (TMAs) were generated in- house from BPPNM tumors harvested 35 days after implantation. TMAs were analyzed by multiplexed immunofluorescence using the PhenoCycler-Fusion system (Akoya Biosciences). Each experimental condition comprised 10 independent tumors, with each tumor represented by two replicate cores containing approximately 1.4 mm² of tissue per core. TMA sections were processed according to the manufacturer’s FFPE protocol and stained with a cocktail of oligonucleotide-barcoded antibodies. Primary antibodies against the following proteins were used: CD11b (E4K8C; CST, 35476SF; 1:100), CD11c (D1V9Y; CST, 39143SF; 1:50), CD8α (D4W2Z; CST, 60168SF; 1:50), F4/80 (D2S9R; CST, 25514SF; 1:100), iNOS (E1W4J; CST, 70706SF; 1:100), Perforin (E3W4I; CST, 36810SF; 1:100), CD19 (D4V4B; CST, 86916SF; 1:50), CD19 (D4V4B; CST, 86916SF; 1:50), Ly6G (E6Z1T; CST, 75028SF; 1:200), CD4 (clone #1; Sino Biological, 50134-R001; 1:50), Pan-Keratin (E6S1S; CST, 72829SF; 1:500), and CD163 (E2H3C; CST, 68922BF; 1:50). Barcode-specific fluorescent reporters conjugated to Atto 550, Alexa Fluor 647, or Alexa Fluor 750 were sequentially hybridized, imaged, and removed, and DAPI was used for nuclear counterstaining. Images were acquired using the PhenoCycler-Fusion system and analyzed with Visiopharm 2024.07 (x64). DAPI-based nuclear segmentation and marker-specific intensity thresholds were used to identify tumor and immune-cell phenotypes. Tissue regions or entire cores affected by tissue loss, folding, staining or imaging artifacts, excessive background, or segmentation failure were excluded before quantification. Spatial relationships were assessed by measuring the distance from each effector T cell (CD8^+^perforin^+^) to the nearest cell of each indicated target population within the same evaluable tissue core. Nearest-neighbor distances were grouped into distance intervals, and the relative frequency (%) in each interval was calculated by dividing the number of effector T cells whose nearest-neighbor distance fell within that interval by the total number of effector T cells included in the corresponding nearest-neighbor analysis for that core and multiplying by 100. Replicate cores were treated as technical replicates, and the individual tumor was used as the biological unit for downstream statistical analysis.

### Human HGSOC sequential immunofluorescence and image analysis

High-grade serous ovarian cancer TMAs were generated in-house and were used for hyperplex sequential immunofluorescence using the Lunaphore COMET. The analyses described here used DAPI together with Pan-cytokeratin (PanCK), CD3, CD4, CD8, granzyme B (GZMB), CD68, CD163, CD11c, and HLA-II. Multichannel images were imported into Visiopharm image-analysis software, version 2024.07 x64 (Visiopharm A/S, Hørsholm, Denmark). Each evaluable TMA core was analyzed as a separate region of interest using a custom, optimized cell-based analysis application. Nuclei were detected using DAPI, and individual cellular objects were segmented and classified according to marker expression. The same optimized analysis application was applied across all evaluable cores. Cells were classified into the following analysis-specific phenotypes: PanCK^+^ epithelial cells, CD8⁺ T cells, CD8⁺GZMB⁺ effector T cells, CD4⁺ T cells, CD11c⁺HLA-II⁺ cells, CD68⁺CD163⁻ tumor-associated macrophages (TAMs), CD68⁺CD163⁺ TAMs, and other cells. Representative pseudocolored cell-classification maps were generated in Visiopharm, with each classified cellular object displayed according to its assigned phenotype. For cell-composition analysis, 21 evaluable TMA cores were included, comprising 14 ancestral *BLTP3A* cores and 7 *BLTP3A* M1098T cores. The single-cell phenotypes were consolidated into four broad categories: T cells, myeloid cells, epithelial cells, and negative/other cells. The T cell category included classified CD8⁺, CD8⁺GZMB⁺, and CD4⁺ T cells. The myeloid category included CD11c⁺HLA- II⁺ cells and the CD68⁺CD163⁻ and CD68⁺CD163⁺ TAM populations. The epithelial category included PanCK⁺ epithelial cells, whereas objects not assigned to these categories were retained as negative/other. The proportion of each category was calculated as a percentage of all classified objects within each core. Core-level cellular compositions were displayed individually and summarized as the mean percentage of classified objects within each genotype group. For spatial analysis, the coordinates of classified cells generated by Visiopharm were used to perform nearest-neighbor analysis. CD8⁺GZMB⁺ cells were designated as effector T cells. The analysis included 256 effector T cells from 14 ancestral *BLTP3A* cores and 231 effector T cells from 7 *BLTP3A* M1098T cores. For each effector T cell, the distance to the nearest PanCK⁺ epithelial cell, CD68⁺CD163⁻ TAM, CD68⁺CD163⁺ TAM, and CD4⁺ T cell was determined. Relative-frequency distributions of effector T-cell distances to each target-cell population were generated over a distance range of 0–100 µm. Spatial proximity between effector T cells and CD68⁺CD163⁻ TAMs was additionally summarized for each core by calculating the area under the distance–frequency curve from 0 to 100 µm^7,70^.

### Immunohistochemistry for BLTP3A

BLTP3A protein expression was assessed by chromogenic immunohistochemistry in formalin-fixed, paraffin-embedded human high-grade serous ovarian carcinoma (HGSOC) tumor specimens. Briefly, 4- to 5-μm tissue sections were baked, deparaffinized in xylene, and rehydrated through graded ethanol to distilled water. Heat-induced epitope retrieval was performed in citrate-based antigen retrieval buffer, pH 6.0, followed by quenching of endogenous peroxidase activity and blocking of nonspecific antibody binding. Sections were incubated with rabbit polyclonal anti-BLTP3A/UHRF1BP1 antibody from the Human Protein Atlas/Atlas Antibodies/Sigma-Aldrich (HPA032099; 1:300) overnight at 4°C. Antibody binding was detected using an HRP-conjugated anti-rabbit polymer detection system and 3,3′- diaminobenzidine (DAB) chromogen, followed by hematoxylin counterstaining, dehydration, clearing, and cover slipping.

### ELISA for conditioned-media cytokines

Secreted cytokines and chemokines in conditioned media were quantified by enzyme-linked immunosorbent assay (ELISA). WT and *Bltp3a*-deficient tumor cells were seeded at equal density (8 × 10⁵ cells per well in 24-well plates) and treated as indicated. Conditioned media were collected 18–24 h after treatment, clarified by centrifugation at 300 × g for 3 min at 4°C to remove cells and debris, and stored at −80°C until analysis.

Mouse CXCL10/IP-10 (PeproTech; #900-K153), IFN-λ2/3 (IL-28A/B; R&D Systems, DuoSet ELISA, DY1789B-05), and CCL5/RANTES (Life Technologies; #88-56009-22) concentrations were measured using commercially available ELISA kits according to the manufacturers’ instructions. Standards and samples were assayed in technical duplicate or triplicate, with three independent biological replicates per condition. Absorbance was measured at 450 nm with wavelength correction at 570 nm. Cytokine concentrations were calculated from plate-specific standard curves following blank subtraction and correction for sample dilution.

### IFN-γ ELISpot assay

IFN-γ ELISpot assays were performed to assess tumor-reactive CD8⁺ T cell responses using precoated mouse IFN-γ ELISpot plates (Cellular Technology Limited [CTL], cat. no. mIFNgp-2M/5). Splenocytes isolated from C57BL/6 mice were pulsed overnight with lysates prepared from BPPNM WT or *Bltp3a*- KO tumors (50 µg/mL). Following antigen loading, splenocytes were irradiated at 32 Gy. Tumor-enriched CD8⁺ T cells (2 × 10⁵ cells per well) were co-cultured with irradiated, lysate-pulsed splenocytes (2 × 10⁴ cells per well) at a 10:1 ratio. Following overnight co-culture, ELISpot plates were developed according to the manufacturer’s instructions, and IFN-γ spot-forming units (SFUs) were quantified by CTL.

### THP-1 macrophage differentiation and conditioned media assay

THP-1 monocytes were differentiated into macrophage-like cells by treatment with 5 ng/mL phorbol 12- myristate 13-acetate (PMA) for 48 h. After differentiation, cells were washed to remove PMA and cultured with conditioned media collected from WT or BLTP3A-deficient OVCAR8 cells. To generate conditioned media, WT and BLTP3A-KO tumor cells were plated at equal density, cultured for 48 h, and supernatants were collected, clarified by centrifugation to remove cells and debris, and applied to differentiated THP- 1 macrophages. THP-1-derived macrophages were then incubated with OVCAR8-conditioned media for 48 h to assess the effect of BLTP3A-dependent tumor-derived factors on macrophage polarization. Macrophage phenotype was evaluated by flow cytometry/qPCR using markers of inflammatory and alternatively activated macrophage states, including CD206, CD163, CXCL9, CD86.

### Immunoprecipitation assay

*BLTP3A*^KO^ OVCAR8 cells were seeded to 60% confluency on 150mm tissue culture and left to adhere overnight. The cells were then co-transfected with Ancestral BLTP3A or M1098T constructs (GenScript) and the EGFP-LC3 construct (Addgene; #11546) using jetPRIME transfection reagent (Sartorius; 101000046) following the manufacturer’s protocol using 20 ug of total DNA. After 24 h, the media was replaced with fresh complete media to reduce cytotoxicity. At 24 h post transfection, some cells were treated with 20% human patient ascites. At approximately 48 h post-transfection, some cells were treated with 3mM L-Leucyl-L-Leucine methyl ester (LLOMe) (Cayman Chemical; #16008) for 1 h. All cells were harvested at 48 h post-transfection via scraping, washed twice with PBS, and pelleted. Subsequent steps were all performed on ice. Pellets were then resuspended in ice-cold IP lysis buffer (Thermo Scientific; #87787) supplemented with 2X protease inhibitor cocktail (Thermo Scientific; #7840) and incubated on ice for 15 minutes with 5-minute intervals of gentle vortex, and then cleared by centrifugation at 13,000 x g for 15 min. The resulting protein lysates were quantified by BCA protein assay (Thermo Scientific; #23225). A portion of the protein lysates were reserved to serve as input controls while equal protein amounts from each experimental condition of the remaining lysates were incubated with anti-GFP magnetic beads (CST; 67090) at 4℃ overnight. Beads were then washed three times with IP lysis buffer, once with PBS, and once with distilled water. The proteins were eluted with 2X LDS sample loading buffer (Thermo Scientific; #NP0007) by heating at 95℃ for 5 min and then de-beaded. Sample reducing agent was then added to the eluent to a final concentration of 1X (Invitogen; #NP0009). 20 ug of input and equal volumes of eluent was ran on a 4-12% Bis-Tris polyacrylamide gel (Invitrogen; #NP0322BOX) and then transferred onto 0.2 µm nitrocellulose membrane (Cytiva; #1600011) using the Xcell Surelock Mini-Cell and blotting module system (Invitrogen; EI0001, #EI9051). Membranes were then blocked with 5% milk and incubated overnight at 4℃ with primary antibodies. Targets include: GFP (CST; #2956), LC3 (CST; #3868), RAB7 (CST; #9367), LAMP1(9091), STING (Proteintech; 19851-1-AP). Membranes were then washed extensively, incubated with secondary HRP conjugated antibody (CST; 7074), and visualized by chemiluminescence.

### SDS-PAGE and immunoblotting

Cells were washed with ice-cold PBS and lysed on ice in RIPA buffer (Millipore; 20-188) supplemented with protease and phosphatase inhibitor cocktails (Thermo Scientific; 78440). Lysates were then cleared by centrifugation at 13,000g/4°C for 15 min, and protein concentrations were determined using a bicinchoninic acid (BCA) assay according to the manufacturer’s instructions (Thermo Scientific; 23225). Equal amounts of protein were mixed with 1X sample LDS loading buffer (Thermo Scientific; NP0007) containing 1X reducing agent (Thermo Scientific; NP0009) and denatured by heating at 70°C for 10 min. Samples were then loaded and ran on 10% or 4-12% Bis-Tris polyacrylamide gels (Invitrogen; NP0315BOX, NP0303BOX, NW04122BOX) and transferred to 0.2µm nitrocellulose membranes (Cytiva; 1600011). Membranes were then blocked for 1 hr at room temperature in 5% non-fat milk (RPI; 50-488-784) or 5% BSA (Sigma-Aldrich; A3912-500G) dissolved in 1X TBST (Sigma-Aldrich; T9039- 10PAK) and incubated overnight at 4°C with primary antibodies diluted in blocking buffer against the indicated proteins. Target proteins include: BLTP3A/UHRF1BP1 (Bethyl Laboratories; A304-646A), STING (Proteintech; 19851-1-AP), phospho-STING (Invitrogen; PA5105674), TBK1 (Invitrogen; 703154), phospho-TBK1 (Invitrogen; MA5-35869), IRF3 (CST; 4920), phospho-IRF3 (Invitrogen; MA5-14947), LC3B (CST; 3868), GABARAP/GABARAPL1 (CST; 26632), TFEB (CST; 32361), LAMP1(CST; 9091), and β-actin (CST; 4970) or GAPDH (CST; 2118) as loading controls, as applicable to each experiment. After washing 3 times with TBST, membranes were incubated with HRP-conjugated secondary antibodies (CST; 7074, 7076) for 1 hr at room temperature, washed 3 times with TBST, incubated with ECL detection reagent (Cytiva; RPN2232), and visualized by chemiluminescence.

### Immunofluorescent Staining and Confocal Microscopy

OVCAR8 or BPPNM cells were seeded on No. 1.5 Poly-D-Lysine coated coverslips to around 50% confluency in a 6-well tissue culture plate (Neuvitro; GG-18-15-PDL). In some experiments after adhering, cells were co-transfected with Ancestral or M1098T FLAG tag BLTP3A constructs (GenScript) and EGFP-LC3 constructs (Addgene; #11546) using jetPRIME® transfection reagent (Sartorius; 101000046) following the manufacturers protocol using 5ug of total DNA. After 24 h, the media was replaced with fresh complete media. Some cells were treated at 24 h post transfection with human or mouse ascites overnight, while others were treated with LLOMe (Cayman Chemical; 16008) for 1 h or 1 h with 3h rest prior to the end point. The cells were kept in culture for a total of 48 h post transfection. Experiments that did not require transfection of constructs were removed from culture when proper confluency was reached and treatments were completed. After removal from culture, coverslip containing cells were gently washed once with PBS in the well and fixed with 4% paraformaldehyde in PBS for 15 min (Thermo; J61899.AK). Cells were then gently washed three times with PBS and blocked for 1 hr with PBS containing 5% normal goat serum (Invitrogen; 10000C) and 0.3% Triton X-100 (Sigma; X100- 500ML). Cells were then incubated at 4℃ overnight with primary antibodies diluted in PBS containing 1% BSA and 0.3% Triton X-100. Targets include FLAG (CST; 14793), STING (Proteintech; 66680-1- Ig), RAB7 (Novus; NBP2-60237), Galectin-3 (Biolegend; 125401-BL). Cells were then gently washed three times with PBS and incubated for 2 h at room temperature with fluorescently tagged secondary antibodies diluted in PBS containing 1% BSA and 0.3% Triton X-100. DAPI was then added to a final concentration of 1uM and incubated for 5 minutes prior to the end of incubation (Invitrogen; D1306). Cells were gently washed three times with PBS and mounted on glass specimen slides with Prolong Diamond mountant (Invitrogen; P36982). Images were acquired with a Nikon A1R-Si confocal microscope using the NIS-Elements software. Colocalization and compartment-specific signals were quantified from identically acquired images using ImageJ.

### Chromatin immunoprecipitation followed by quantitative PCR

Candidate ATF4-binding motifs within regulatory regions of *BLTP3A/UHRF1BP1* were identified using JASPAR and visualized in the UCSC Genome Browser using the GRCh37/hg19 human genome assembly. OVCAR3 cells were treated with FCCP (1 μM) or DMSO vehicle for 16 h. Cells were crosslinked with 1% formaldehyde for 10 min at room temperature, and crosslinking was quenched with 125 mM glycine for 5 min. Cells were washed with ice-cold PBS, collected, and lysed in the presence of protease inhibitors. Chromatin was sheared to an average fragment size of approximately 200–500 bp using a Bioruptor Pico (Diagenode). An aliquot corresponding to 2.5% of each chromatin preparation was retained as input. Equivalent amounts of sheared chromatin were immunoprecipitated with ATF-4 (D4B8) Rabbit monoclonal antibody (Cell Signaling Technology, #11815) or Normal Rabbit IgG (Cell Signaling Technology, #2729) using ChIP-grade Protein G Magnetic Beads (Cell Signaling Technology, #9006S). Following immunoprecipitation, beads were washed and bound chromatin was eluted. Crosslinks were reversed, proteins were digested with proteinase K, and DNA was purified using the QIAquick PCR Purification Kit (QIAGEN). Immunoprecipitated and input DNA were analyzed by quantitative PCR using Luna Universal qPCR Master Mix (New England Biolabs). qPCR reactions were performed in quadruplicate on a CFX384 Real-Time PCR Detection System (Bio-Rad). Primer pairs amplified candidate ATF4-binding regions located approximately 1.3 and 3.4 kb upstream of the annotated *BLTP3A/UHRF1BP1* transcription start site. ChIP enrichment was calculated as percent input after accounting for the 2.5% input fraction, with rabbit IgG immunoprecipitation used to assess nonspecific chromatin recovery^71^.

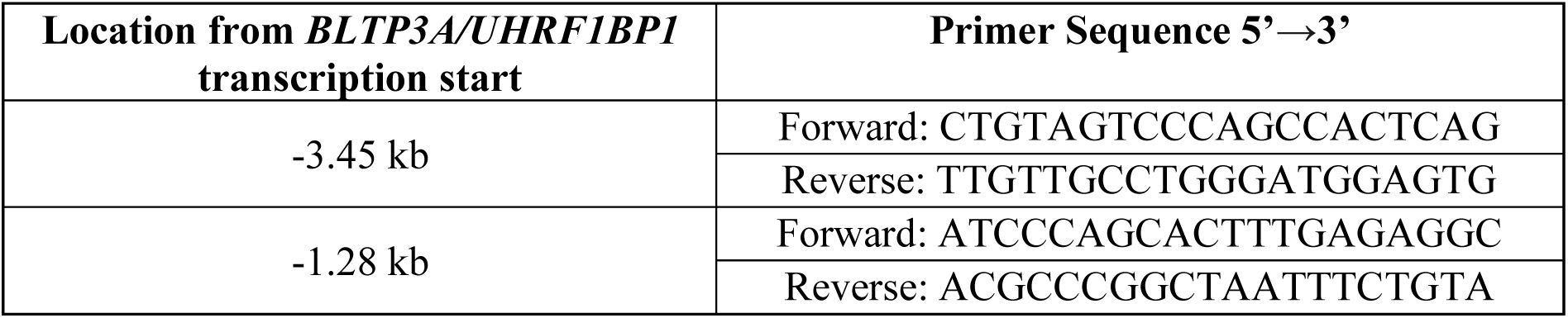

### Immunoprecipitation and LC-MS/MS Analysis

For interactome analysis, *BLTP3A*-deficient OVCAR3 cells were reconstituted with C-terminal FLAG- tagged ancestral *BLTP3A*, *BLTP3A*^M1098T^, or empty-vector control and treated with FCCP or DMSO vehicle as indicated. Where indicated, intracellular protein complexes were stabilized before lysis by incubation with 2 mM dithiobis (succinimidyl propionate) (DSP; Lomant’s reagent) for 30 min at room temperature, followed by quenching with Tris. Cells were washed extensively with ice-cold PBS, harvested, and lysed on ice in immunoprecipitation lysis buffer supplemented with protease inhibitors. Lysates were clarified by centrifugation, and protein concentrations were determined using a BCA protein assay. An aliquot of each lysate was retained as an input control, and equivalent amounts of protein from the remaining lysates were subjected to FLAG immunoprecipitation using Pierce DYKDDDDK Magnetic Agarose (Thermo Fisher Scientific) at 4°C. Beads were washed extensively to remove nonspecifically associated proteins, and FLAG-associated protein complexes were eluted in LDS sample buffer.

For LC–MS/MS analysis, eluates were briefly resolved by SDS-PAGE and visualized by colloidal Coomassie staining. Protein-containing gel regions were excised and subjected to in-gel proteolytic digestion before LC–MS/MS analysis at the Rutgers Center for Advanced Proteomics Research.

Candidate BLTP3A-interacting proteins were evaluated relative to the corresponding empty-vector immunoprecipitation controls and compared across BLTP3A genotype and treatment conditions. BLTP3A itself was excluded from downstream interactor analyses. Protein enrichment was calculated relative to matched empty-vector controls, and candidate interactors were prioritized based on reproducible enrichment across biological replicates and statistical support. Where indicated, interaction enrichment was expressed as the log2-transformed ratio of protein abundance in BLTP3A-FLAG immunoprecipitates relative to the corresponding empty-vector control.

### Murine Single-cell RNA-sequencing analysis

CD45⁺ immune cells were isolated on day 35 after tumor inoculation from mice bearing WT BPPNM tumors (n = 5) or *Bltp3a*KO BPPNM tumors (n = 3). Tissues were dissociated as described in the Tumor dissociation section, and viable CD45⁺ singlets were isolated by fluorescence-activated cell sorting (FACS). Samples from individual mice were processed separately.

Single-cell gene-expression libraries were generated using the Chromium Next GEM Single Cell 5′ Gene Expression platform by Novogene (10x Genomics). Matched T cell receptor (TCR) V(D)J libraries were generated from the same cell preparations using the Chromium Single Cell Mouse T Cell V(D)J Enrichment Kit according to the manufacturer’s instructions. Gene-expression and V(D)J libraries were sequenced on an Illumina NovaSeq X Plus platform using paired-end, dual-indexed sequencing with 26 cycles for Read 1, 10 cycles each for the i7 and i5 index reads, and 90 cycles for Read 2, targeting a minimum depth of 20,000 read pairs per cell for gene-expression libraries and 5,000 read pairs per cell for V(D)J libraries. Demultiplexed FASTQ files were processed using Cell Ranger version 8.0.1 (10x Genomics), with the cellranger count pipeline used for gene-expression libraries and the cellranger vdj pipeline used for TCR V(D)J libraries. Gene-expression reads were aligned to the 10x Genomics mouse reference transcriptome refdata-gex-GRCm39-2024-A (GRCm39; GENCODE vM33/Ensembl 110), and cell barcodes and unique molecular identifiers (UMIs) were used to generate gene-by-cell count matrices. V(D)J reads were assembled and annotated using the 10x Genomics mouse V(D)J reference refdata- cellranger-vdj-GRCm38-alts-ensembl-7.0.0 to reconstruct full-length, productive TCR α- and β-chain sequences and assign clonotypes.

Gene-by-cell UMI count matrices were analyzed using Piccolo^72^ (version 1.0.1). Quality-control filtering was performed independently for each sample. Cells expressing fewer than 200 genes, cells with mitochondrial transcripts accounting for more than 30% of total UMIs, and cells with total UMI counts greater than the sample median plus 3.5 median absolute deviations were excluded. Genes detected in fewer than 0.5% of cells passing quality-control filtering were removed. We used *scDblFinder* with default parameters^73^ (version 1.26.7) for identifying doublets, and no cells were removed from the downstream analysis. Following quality control, 65,039 cells were retained, including 34,641 cells from WT samples and 30,398 cells from KO samples. A k-nearest-neighbor graph was constructed using the first 50 principal components, with k = 10. Cell clusters were identified using Leiden clustering method at a resolution of 1.0. Uniform Manifold Approximation and Projection (UMAP) was used for visualization. Cluster separability was assessed using cluster-specific marker-gene sets. For each cluster, marker-gene expression was used to calculate a composite z score for every cell. The ability of the composite score to distinguish cells within the corresponding cluster from all other cells was evaluated using the area under the precision-recall curve (AUPRC) and area under the receiver operating characteristic curve (AUROC). Clusters with an AUPRC ≥ 0.625 and an AUROC ≥ 0.85 were retained, resulting in 27 final clusters. Candidate cell identities were assigned by comparing cluster-enriched genes with curated mouse cell-type markers in CellMarker 2.0. Annotations were subsequently refined by manual inspection of canonical lineage markers, cluster-specific marker genes, and broader transcriptional profiles. Cells annotated as T cells or monocytes/macrophages were extracted from the combined dataset and independently reanalyzed using Piccolo. Subclusters were identified using Louvain clustering method at a resolution of 1.0, resulting in 17 T-cell subclusters and 13 monocyte/macrophage subclusters. Subcluster identities were assigned by manual evaluation of canonical lineage, differentiation, activation, and functional-state markers. Differences in cell-population abundance between WT and KO tumors were evaluated using a two-sided Mann–Whitney U test, with each mouse treated as an independent biological replicate. P values were adjusted for multiple comparisons using the Benjamini–Hochberg method. Differential gene-expression analyses between WT and KO cells were performed separately within individual cell-type populations using Piccolo. Genes with an adjusted p value < 0.05 and an absolute log₂ fold change greater than 1 were considered differentially expressed. Individual mice were treated as independent biological replicates, and sample identity was incorporated into differential-expression analysis. Pseudotime trajectory analysis was performed on monocyte and macrophage populations from *Bltp3a*^KO^ tumors using Monocle 3 version 1.4.2. Cells of interest were extracted from the Piccolo object, and the corresponding raw UMI counts and cell-level metadata were converted into a Monocle 3 cell_data_set object. A principal trajectory graph was inferred using learn_graph with the default parameters on the monocyte/macrophage subset of cells on the pre-analyzed dimension. The trajectory root was assigned to the transcriptionally defined monocyte population based on expression of canonical monocyte markers, and cells were ordered along the inferred trajectory using order_cells.

### Single-cell TCR V(D)J repertoire analysis

High-confidence, full-length, productive TCR contigs identified by Cell Ranger were retained for repertoire analysis. V(D)J sequences were linked to transcriptomic cell identities using shared cell barcodes. A clonotype was defined based on the Cell Ranger clonotype definition, a group of adaptive T cells that share a common fully recombined V(D)J ancestor. Clonal expansion was quantified from the frequency of each clonotype within individual biological samples and annotated T cell populations.

TCR repertoire diversity was quantified using the inverse Simpson index. To account for differences in T cell recovery among samples, repertoires were rarefied to a common number of productive paired TCRαβ cells before calculation of diversity metrics. Diversity and clonal-expansion metrics were compared between genotypes using a two-sided Mann–Whitney U test, with individual mice treated as independent biological replicates. Productive CDR3β amino acid sequences were analyzed using GLIPH2^35^ to identify putative TCR convergence groups. GLIPH2 groups TCR sequences based on enriched local CDR3 motifs and global sequence similarity and therefore predicts shared antigen-recognition features rather than demonstrating common antigen specificity. GLIPH2 was run on the combined samples using the mouse reference repertoire presented by the website. Convergence groups were retained when they met the following prespecified criteria significant V-gene enrichment (vb_score ≤ 0.05), significant motif/cluster enrichment based on Fisher’s exact test (Fisher_score ≤ 0.05), a stringent overall convergence score (−log(final_score) > 18, and motif pattern length > =4). Paired αβ clonotype frequencies were calculated as the number of TCR-matched cells belonging to each clonotype divided by the total number of TCR- matched cells within the corresponding biological sample. For expanded clonotypes, distribution across transcriptional T cell subpopulations was calculated as the fraction of cells from each clonotype assigned to each T cell subpopulation.

### Bulk RNA sequencing and differential expression analysis

Bulk RNA sequencing was performed using independent biological replicates of WT and KO BPPNM cells cultured under untreated conditions or exposed to 20% (v/v) cell-free ovarian cancer ascites for 18 h. The experimental design therefore comprised four groups: WT untreated, WT plus ascites, KO untreated, and KO plus ascites, with 3 independent biological replicates per condition. Total RNA was isolated using the RNeasy Mini Kit (QIAGEN) according to the manufacturer’s instructions. Poly(A)⁺ mRNA was enriched from total RNA using oligo(dT)-conjugated magnetic beads, and strand-specific RNA-seq libraries were prepared by Novogene according to its standard library-preparation workflow. Libraries were sequenced on an Illumina NovaSeq X Plus platform using 150-bp paired-end reads to a depth of approximately 20 million read pairs, corresponding to approximately 6 Gb of raw sequence data per sample. Transcript abundance was quantified directly from FASTQ files using kallisto v0.46.1, with a transcriptome index generated from the *Mus musculus* Ensembl v96 annotation. Differential gene- expression analysis was performed in R version 4.5.2 using DESeq2 version 1.50.2. Genes were retained when they had 10 in at least 2 samples. Differential expressions were evaluated using the Wald test. P values were adjusted for multiple testing using the Benjamini–Hochberg false-discovery rate procedure. Genes with an adjusted p value < 0.05 and an absolute log₂ fold change greater than 1 were considered differentially expressed. Gene-set enrichment analysis was performed using fgsea version 1.36.2 in R. For each reported comparison, genes were ranked according to the DESeq2 Wald statistic. Enrichment analysis was performed using Molecular Signatures Database gene sets, including the Hallmark, Gene Ontology Biological Process, and KEGG collections. Gene sets with an FDR < 0.25 and a nominal p value < 0.05 were considered enriched. Normalized enrichment scores were used to indicate the direction and magnitude of pathway enrichment.

### Visium HD spatial transcriptomic analysis

Formalin-fixed, paraffin-embedded (FFPE) high-grade serous ovarian carcinoma (HGSOC) specimens were processed for Visium HD Spatial Gene Expression analysis (10x Genomics^74^). Tissue sections were mounted onto glass slides and processed according to the manufacturer’s Visium HD FFPE and CytAssist protocols. Following tissue pretreatment and decrosslinking, whole-transcriptome oligonucleotide probes were hybridized to tissue RNA and ligated. The Visium CytAssist instrument was used to transfer ligated probe products onto spatially barcoded Visium HD capture slides. Following probe capture and extension, sequencing libraries were amplified, indexed, quality-controlled, and sequenced.

Visium HD spatial transcriptomic data from 13 HGSOC specimens were analyzed at 8-µm resolution to investigate associations between epithelial lysosomal remodeling and local immune organization. Sequencing data were processed using Space Ranger v4.0.1 (10x Genomics), the GRCh38-2024-A human reference, and Visium Human Transcriptome Probe Set v2.1.0. Specimen-matched CytAssist TIFF images were used for automated image registration and tissue identification; tissue detection was enabled and cell segmentation was disabled. Filtered 8-µm count matrices, spatial coordinates, and images were imported into Seurat using the *Spatial.008um* assay. During object reconstruction, zero-count regions, duplicate or malformed barcodes, and regions lacking valid spatial coordinates were removed. Raw transcript counts were retained for downstream analyses.

Epithelial-enriched regions were identified using high-specificity lineage-anchor annotations established through marker-based and reference-guided compartment classification. An initial sketch of 350,000 regions from 14 specimens was evaluated using epithelial, immune, stromal, and endothelial marker signatures and donor-balanced HGSOC single-cell transcriptomic reference profiles. Reference-guided scores combined expression-correlation probabilities with nonnegative mixture-decomposition estimates. The final epithelial marker panel comprised *EPCAM, KRT8, KRT19, KRT7, MSLN, MUC1, WFDC2,* and *KRT17*. Epithelial-anchor assignment required the epithelial compartment to have the highest consensus score, a consensus probability ≥0.50, at least 11 detected transcripts and 11 detected genes, and a calibrated epithelial marker-contrast threshold ≥−1. Thresholds were calibrated using donor-held-out synthetic spatial mixtures. Mixed and ambiguous regions were excluded. The resulting epithelial-enriched regions were used for subsequent spatial analyses.

Epithelial lysosomal remodeling was assessed using a composite transcriptional score encompassing five programs associated with lysosomal homeostasis: lysosomal biogenesis, degradation, damage/repair, membrane protection, and lysophagy^75–80^. The 62-gene composite signature comprised *ACP2, ANXA1, ANXA2, ANXA6, ASAH1, ATG5, ATG7, ATP13A2, ATP6V0C, ATP6V0D1, ATP6V1A, ATP6V1B2, ATP6V1H, BECN1, CALCOCO2, CHMP2A, CHMP2B, CHMP4B, CHMP7, CLN3, CTSB, CTSD, CTSL, CTSS, CTSZ, DNASE2, GAA, GABARAPL2, GLB1, GUSB, HEXA, HEXB, HSPA1A, HSPA8, LAMP1, LAMP2, LGALS3, LGALS8, LIPA, MAN2B1, MAP1LC3B, MCOLN1, MITF, NAGA, NPC2, OPTN, PDCD6IP, PPT1, PSAP, RAB7A, RUBCN, SCARB2, SORT1, SQSTM1, TAX1BP1, TBK1, TFE3, TFEB, TSG101, VPS4A, VPS4B,* and *WIPI2*. Transcript counts were normalized to 10,000 transcripts per region and log-transformed. For each lysosomal component, individual gene-expression values were ranked across eligible epithelial regions within each specimen, and normalized ranks were averaged across genes in the corresponding signature. Component scores were standardized within each specimen and combined using an equal-weight mean to generate the composite lysosomal-remodeling score. Within each specimen, epithelial-enriched regions in the upper and lower quartiles of the composite score were designated lysosomal-remodeling-high and -low, respectively. Intermediate regions were excluded from comparisons.

Myeloid/macrophage-lineage regions were identified using a gene-expression signature comprising *AIF1, CD68, CSF1R, CTSS, FCER1G, LST1, LYZ, MAFB, MS4A7, SPI1,* and *TYROBP*. Classification required detection of at least two signature genes and a lineage score within the upper quartile for each specimen. Within these regions, *CXCL9*^+^ inflammatory antigen-presenting-cell (APC)-like myeloid states were evaluated using a 12-gene signature comprising *CD40, CD80, CXCL9, CXCL10, CXCL11, GBP1, GBP5, IDO1, IRF1, STAT1, TAP1,* and *TAP2*. This signature encompasses inflammatory chemokine production, interferon responsiveness, and antigen-presentation-associated transcriptional programs characteristic of immunostimulatory myeloid populations^7,30^. Myeloid program scores were calculated from normalized gene expression using positive-expression ranks. Local APC-like myeloid enrichment was evaluated within an 80-µm radius of lysosomal-remodeling-high and -low epithelial regions.

To assess the spatial relationship between APC-like myeloid populations and T cells, we quantified T cell- associated regional abundance within 80 µm of APC-like-positive and APC-like-negative myeloid regions. T cell-enriched regions were identified at 16-µm resolution using a lineage signature comprising *CD2, CD3D, CD3E, CD3G, CD247, TRAC, TRBC1,* and *TRBC2*. Classification required detection of at least two signature genes and a lineage score within the upper quartile for each specimen. Local T cell abundance was estimated from the number of T cell-signature-positive 16-µm regions within each neighborhood. These counts were log-transformed and standardized within specimens for spatial comparisons.

To determine whether epithelial transcriptional states were spatially associated with immune cell populations beyond what would be expected from local tissue organization and technical variation, observed spatial associations were adjusted against a matched null model. Specifically, for each tumor specimen 100 random draws were generated, matching observations within 200-µm spatial blocks by sequencing-depth and detected-feature-count quartiles. The mean difference across these matched random draws was used to estimate the expected spatial difference under the null model. The specimen-level adjusted effect was then calculated by subtracting this mean matched-null difference from the observed difference between lysosomal-high and -low regions, or between APC-like-positive and negative regions. Cohort-level effects were summarized using Hodges–Lehmann location estimates and Wilcoxon signed- rank confidence intervals, where available^81^. Two-sided Wilcoxon signed-rank tests were used to assess statistical significance across independent tumor specimens, using one adjusted effect estimate per specimen. Multiple comparisons were corrected using the Benjamini–Hochberg procedure, applied separately to the epithelial–myeloid and myeloid–T cell spatial analyses^82^. Findings were interpreted as spatial associations.

### TCGA rs13205210 survival analysis

Overall survival was evaluated in The Cancer Genome Atlas ovarian cancer cohort (TCGA-OV) among participants with callable germline rs13205210 genotypes derived from matched-normal whole-exome sequencing. Participants were classified as ancestral-genotype homozygotes (TT; n = 243) or M1098T carriers, with heterozygous and homozygous carriers combined (CT/CC; n = 76). Associations with overall survival were estimated using Cox proportional hazards regression adjusted for age at diagnosis, disease stage, and the first five genetic-ancestry principal components. Kaplan–Meier curves were used for visualization, and unadjusted group comparisons were performed using the log-rank test.

### Human rs13205210 genotyping

Genomic DNA was isolated from human HGSOC tumor specimens using a QIAGEN DNA extraction kit according to the manufacturer’s instructions. DNA concentration and purity were assessed using a NanoDrop spectrophotometer (Thermo Fisher Scientific). The BLTP3A/UHRF1BP1 M1098T variant rs13205210 was genotyped using a TaqMan SNP Genotyping Assay (Applied Biosystems/Thermo Fisher Scientific) with TaqMan Genotyping Master Mix. Each 10 μL reaction contained 10 ng genomic DNA, TaqMan Genotyping Master Mix, SNP-specific assay mix containing allele-specific VIC- and FAM- labeled probes, and nuclease-free water. Amplification and endpoint fluorescence detection were performed on an Applied Biosystems StepOnePlus Real-Time PCR System using the manufacturer- recommended cycling conditions. Genotypes were assigned by allelic discrimination using StepOne Software v2.3. No-template controls and, when available, DNA samples of known genotypes were included. Samples with failed amplification, ambiguous allelic clustering, insufficient DNA quality, or discordant replicate genotype calls were excluded from downstream analyses.

### *BLTP3A* expression in malignant and nonmalignant human ovarian cancer epithelial cells

To evaluate BLTP3A expression across malignant and nonmalignant epithelial populations, we analyzed the concatenated human ovarian cancer single-cell RNA-sequencing dataset. Cells annotated as epithelial/tumor cells were identified using the major cell-type annotations and classified as malignant or nonmalignant according to available pathology, disease-stage, and sample annotations. The nonmalignant group included epithelial cells from normal-adjacent tissue (NAT) and benign or non-cancer specimens. BLTP3A transcript abundance was extracted from the normalized RNA expression matrix using its annotated gene symbol, *UHRF1BP1*. For each patient, mean normalized BLTP3A expression, median expression, and the fraction of cells with detectable BLTP3A transcripts (normalized expression greater than zero) were calculated. Patient-level mean BLTP3A expression was compared between malignant and nonmalignant epithelial populations using a two-sided Wilcoxon rank-sum test, with patients serving as the unit of statistical analysis.

### Transcriptional profiling of lysosomal stress and STING signaling in human ovarian cancer

To characterize lysosomal stress responses and their association with STING signaling in human high- grade serous ovarian carcinoma (HGSOC), we analyzed malignant epithelial cells from our previously assembled, concatenated human ovarian cancer single-cell RNA-sequencing dataset. Normalized expression values were standardized on a per-gene basis, and transcriptional program scores were calculated as the unweighted mean of standardized expression values for genes represented in the dataset. Lysosomal stress responses were evaluated using distinct transcriptional programs representing lysophagy and lysosomal membrane protection. The lysophagy program was defined by established mechanisms of selective autophagic recognition and clearance of damaged lysosomes^46,77^and comprised *LGALS3, LGALS8, CALCOCO2, SQSTM1, OPTN, TAX1BP1, TBK1, ATG5, ATG7, BECN1, MAP1LC3B, GABARAPL2, WIPI2, LAMP2, RAB7A,* and *RUBCN*. The lysosomal membrane-repair program represented proteins associated with lysosomal membrane integrity, homeostasis, and stress adaptation and comprised *LAMP1, LAMP2, SCARB2, MCOLN1, ATP13A2, HSPA1A,* and *HSPA8*.

The productive STING-output signature was designed to capture downstream interferon-responsive transcription and comprised *ISG15, IFIT1, IFIT2, IFIT3, IFI6, IFI44, IFI44L, MX1, OAS1, OAS2, OAS3, CXCL10, CCL5, STAT1, STAT2, IRF7, DDX58,* and *IFIH1*. Lysophagy and membrane-repair scores were independently correlated with productive STING-output scores using Spearman rank correlation at the patient level. The primary analysis included 48 independent patients with malignant HGSOC epithelial cells. Multiple comparisons were adjusted using the Benjamini–Hochberg procedure.

### Statistical analysis

Statistical analyses were performed in R and GraphPad Prism. Two-sided tests were used unless otherwise stated. Wilcoxon rank-sum tests were used for two-group comparisons, Kruskal–Wallis tests for multigroup comparisons, and Spearman rank correlations for associations between gene programs. Linear regression lines in scatter plots were included for visualization unless the corresponding regression model was explicitly specified. Multiple-testing correction used the Benjamini–Hochberg procedure within prespecified analysis families. Survival was analyzed using Kaplan–Meier estimation, log-rank testing, and covariate-adjusted Cox regression. The biological replicate was the mouse, human participant, tumor, or independently generated culture, as appropriate; technical replicates were not treated as independent.

## ACKNOWLEDGEMENTS

This study was supported by NCI R37CA295820 and L30CA284429, the Pershing Square Sohn Cancer Prize, the V Foundation for Cancer Research (AST2025-011 and V2022-030), the Ovarian Cancer Research Alliance (ECIG-2023-3-1007), and Rutgers Cancer Institute start-up funds to K.K. Payne. E.S. Hosseini was supported by NCI 1F99CA305528-01; M. Arabzadeh, R. Kumari, C. Alcott, and A. Robinson by New Jersey Commission on Cancer Research awards COCR24PDF008, COCR23PDF002, COCR26PRF004, and COCR27PPR018, respectively; R. Kumari by American Cancer Society fellowship PF-23-1142024-01-IBCD; and A. Robinson by NIH T32GM139804-05. This work was supported in part by Rutgers Cancer Institute CCSG P30CA072720 and its Biomedical Informatics, Immune Monitoring and Flow Cytometry, and Biospecimen Repository and Histopathology Shared Resources. Additional support was provided by the Cellular Imaging and Histology Core and Center for Advanced Metabolomics and Proteomics Research at Rutgers New Jersey Medical School, and the Immune Monitoring and Cancer Omics Services at OHSU Knight Cancer Institute. We thank Ankit Saxena, Kelly Walton, Joseph Rosenberg, Nikolaus Svoronos, Nadia Kim, Sanjana Philips, and Tyler Milonas for technical assistance; Dr. Yonglun Luo (Aarhus University) and Dr. Robert Weinberg (Whitehead Institute) for providing the ovarian cancer single-cell dataset and BPPNM models, respectively; and Dr. Barth Grant (Rutgers University) for helpful discussions. We are especially grateful to Dr. José R. Conejo-Garcia (Duke University) for his continued guidance and support.

