## Supplemental Figures and Legends for "BLTP3A dysfunction unleashes lysosomal stress–induced antitumor immunity through STING"

#### SUPPLEMENTAL FIGURE LEGENDS

**Figure S1. Human and experimental validation of BLTP3A-associated ovarian cancer control. (A)** Representative multiplex immunofluorescence images of $BLTP3A^{\mathrm{Ancs}}$ and $BLTP3A^{\mathrm{M1098T}}$ HGSOCs showing CD3, GzmB, Pan-CK, and DAPI. **(B)** $CD3^{+}GzmB^{+}$effector T cell frequency among classified cells in 22 ancestral, 7 heterozygous M1098T, and 6 homozygous M1098T tumors (n=22, n=7, and n=6). Cellular composition of individual tumor cores analyzed using Visiopharm Phenoplex software. **(C)** Stacked bars show the percentages of T cells, myeloid cells, epithelial cells, and negative or unclassified objects in $BLTP3A^{\mathrm{Ancs}}$ and $BLTP3A^{\mathrm{M1098T}}$ specimens. **(D)** Mean cellular composition of $BLTP3A^{\mathrm{Ancs}}$ and $BLTP3A^{\mathrm{M1098T}}$ tumors analyzed in (C). **(E)** Patient-level mean of *BLTP3A* expression in nonmalignant and malignant epithelial cells from the integrated ovarian cancer single-cell RNA-sequencing atlas. **(F)** BLTP3A protein abundance in $BLTP3A^{\mathrm{Ancs}}$ and $BLTP3A^{\mathrm{M1098T}}$ human tumors, quantified by β-actin-normalized immunoblot densitometry. **(G)** Longitudinal tumor bioluminescence in mice bearing luciferase-expressing WT or $Bltp3a^{\mathrm{KO}}$ BPPNM tumors. **(H)** Tumor weights measured at a matched time point (day 35) in mice bearing WT or $Bltp3a^{\mathrm{KO}}$ BPPNM tumors. **(I)** Tumor-rechallenge design, representative bioluminescence images, and longitudinal tumor signal. Mice surviving more than 90 days after rejection of 3 x $10^6$ $Bltp3a^{\mathrm{KO}}$ BPPNM cells were rechallenged intrapretoneally with 3 x $10^6$ WT BPPNM cells; naive mice challenged in parallel served as controls. Tumor bioluminescence was monitored for 33 days (Control, n= 3; rechallenge, n= 3). **(J)** Survival of immunodeficient mice challenged with WT or $Bltp3a^{\mathrm{KO}}$ BPPNM cells (n= 5 mice per group). **(K)** Experimental design for conditional *Kras* activation and *Trp53*/*Bltp3a* deletion in the orthotopic ovarian model and survival of *Bltp3*a-intact and *Bltp3a*-deficient cohorts. **(L)** Cross-species alignment of the BLTP3A amino acid sequence surrounding human M1098 and orthologous murine M1090. Data are mean ± SEM unless otherwise indicated;

Survival curves were compared by log-rank tests; other comparisons used the tests described in Methods. *$p < 0.05$, **$p < 0.01$, ***$p < 0.001$; ns, not significant.

**Figure S2. *Bltp3a* loss remodels tumor-infiltrating T cell states, spatial organization, and clonality.** **(A)** Dot plot of canonical lineage and cell-state markers across intratumoral macrophage/monocyte, dendritic cell, neutrophil, NK cell, T cell, and B cell populations. **(B)** Genotype-separated UMAPs of transcriptionally defined T cell states from WT and *Bltp3a*$^{KO}$ tumors. **(C)** Dot plot of genes associated with lineage identity, activation, exhaustion, regulatory T cell programs, interferon responses, and memory across transcriptionally defined T cell states. **(D)** Relative frequencies of the indicated T cell states in WT and *Bltp3a*$^{KO}$ tumors. **(E)** Feature plots of *Gzmk*, *Ctla4*, and *Sell* expression across the T cell UMAP. **(F)** Dot plot of selected activation- and inhibitory-receptor genes in WT and *Bltp3a*$^{KO}$ CD8$^{+}$ T cells. **(G)** Area under the 0–100-μm distance-frequency curves for effector T cells neighboring iNOS$^{+}$CD163$^{-}$ TAMs in WT and *Bltp3a*$^{KO}$ tumors. **(H)** Representative flow-cytometry of GzmB expression in intratumoral CD8$^{+}$ T cells from WT and Bltp*3a*$^{KO}$ tumors. **(I)** GLIPH2-defined CDR3β motifs unique to or shared between WT and *Bltp3a*$^{KO}$ tumor T cell repertoires. **(J)** Inverse Simpson diversity index of the recovered intratumoral TCR repertoire. **(K)** Representative flow-cytometric validation of CD8$^{+}$ T cell depletion in tumor-bearing mice treated with anti-CD8 or IgG isotype control. Data are mean ± SEM unless otherwise indicated. *$p < 0.05$, **$p < 0.01$, ***$p < 0.001$; ns, not significant.

**Figure S3. Functional characterization of the inflammatory myeloid compartment in *Bltp3a*-deficient tumors.** **(A)** Dot plot of canonical marker genes across *C1q*$^{+}$, *Cxcl9*$^{+}$, *Spp1*$^{+}$, *Isg15*$^{+}$, and *Ddit4*$^{+}$ stress-response TAMs, *Il1b*$^{+}$ inflammatory macrophage, *Ccr2*$^{+}$ inflammatory-monocyte, PMN-MDSC-like cells. **(B)** Feature plots of *Spp1* and *Cxcl9* expression across the macrophage–monocyte UMAP. **(C)** Flow-cytometric gating strategy for viable CD45$^{+}$CD11b$^{+}$ tumor-infiltrating myeloid cells, Ly6C$^{hi}$Ly6G$^{-}$ monocytes, neutrophil/PMN populations, and F4/80$^{+}$MHC-II$^{+}$ TAMs. **(D)** Representative histograms and quantification of tumor-derived GFP signal in CD11b$^{+}$F4/80$^{+}$MHC-II$^{+}$ macrophages from WT and *Bltp3a*$^{KO}$ tumors. **(E)** Experimental design for implantation of H-2K$^{b}$-expressing *Bltp3a*$^{KO}$ BPPNM cells into H-2K$^{d}$ BALB/c hosts under CD8-depleting conditions and quantification of tumor-derived H-2K$^{b}$ acquired by CD86$^{+}$CD206$^{-}$ and CD86$^{-}$CD206$^{+}$ host macrophages. **(F)** Representative flow cytometric assessment of intratumoral macrophage depletion following anti-Gr-1 treatment. **(G)** Frequency of CD8$^{+}$ TILs in WT and *Bltp3a*$^{KO}$ tumors following isotype or anti-Gr-1 treatment. **(H)** Representative SIINFEKL-tetramer and Ki-67 flow cytometry from OT-I adaptive transfer experiments, with or without FTY720. Data are mean ± SEM unless otherwise indicated. *$p < 0.05$, **$p < 0.01$, ***$p < 0.001$; ns, not significant.

**Figure S4. Tumor-derived IFN-λ2/3 is required for *Bltp3a*-deficient tumor control and is therapeutically actionable.** **(A)** Representative CD206 and MHC-II flow cytometry of TAMs isolated from *Rag2*$^{-/-}$ mice bearing WT or *Bltp3a*$^{KO}$ tumors. **(B)** Representative flow cytometry of MHC-II$^{+}$F4/80$^{+}$ TAMs and CD8$^{+}$ and CD4$^{+}$ T cells in *Bltp3a*$^{KO}$ tumors with or without deletion of *Ifnl2/3*$^{KO}$. **(C)** Survival of mice bearing *Bltp3a*$^{KO}$ tumors treated with anti-IFNAR1 or IgG isotype control. **(D)** Survival of mice bearing WT tumors treated with mock plasmid or a DNA-encoded, tumor-targeted IFN-λ procytokine. **(E)** CD163 expression in PMA-differentiated THP-1 macrophages treated with recombinant IFN-λ at the indicated concentrations. **(F)** Representative flow cytometry of human CD45 (hCD45) reconstitution in peripheral blood and spleen of HLA-A2-matched, cord-blood-humanized NSG-SGM3 mice. **(G)** Frequency of tumor-infiltrating human CD45$^{+}$ (hCD45$^{+}$) leukocytes among live cells in tumors of indicated OVCAR3 genotypes**.** **(H)** Representative flow-cytometry plots and quantification of human CD68$^{+}$CD163$^{+}$ macrophages recovered from WT and *BLTP3A*-deficient OVCAR3 tumors in humanized mice. Data are mean ± SEM unless otherwise indicated; Survival curves were compared by log-rank tests;

other comparisons used the tests described in Methods. *$p < 0.05$, **$p < 0.01$, ***$p < 0.001$; ns, not significant.

**Figure S5. BLTP3A recruitment and induction within a lysosomal stress–STING response network.** **(A)** Selected pathways enriched in cancer cells following ovarian cancer ascites exposure by ranked gene set enrichment analysis. **(B)** Representative confocal images and quantification of Gal-3 puncta in WT and *Bltp3a*$^{KO}$ BPPNM cells following LLOMe treatment. **(C)** Local enrichment of CD8$^{+}$ T cell-associated regions near myeloid regions with a CXCL9/CXCL10-associated APC-like signature across 13 HGSOC samples. Circles represent individual samples, and diamonds denote the cohort's summary. Cohort-level q = 0.0168 **(D)** Heatmap of selected proteins enriched by BLTP3A$^{OE}$ or BLTP3A$^{M1098T}$ immunoprecipitates relative to mock control. **(E)** Representative confocal images of BLTP3A and RAB7 in OVCAR8 cells treated with vehicle or FCCP. **(F)** STING immunoblot time course in WT and *Bltp3a*$^{KO}$ cells exposed to 20% ovarian cancer ascites for the indicated times. **(G)** STING protein abundance following treatment with the direct STING agonist diABZI in WT and *Bltp3a*$^{KO}$ BPPNM cells. (**H**) Relative *BLTP3A* expression in OVCAR8 cells treated with vehicle, LLOMe, or LLOMe plus ISRIB. **(I)** Relative *Bltp3a* expression in WT and *Sting1*$^{KO}$ BPPNM cells following vehicle, LLOMe, or 20 % ascites exposure. Spearman correlation coefficients (ρ) and corresponding *P* values are indicated. Data are mean ± SEM unless otherwise indicated; other comparisons used the tests described in Methods. *$p < 0.05$, **$p < 0.01$, ***$p < 0.001$; ns, not significant.

**Figure S6. *Bltp3a* loss alters STING compartmentalization without detectable cGAMP elevation.** **(A)** Intracellular cGAMP concentrations in WT and *Bltp3a*$^{KO}$ BPPNM cells treated with vehicle, 20% ovarian cancer ascites, or 1 mM LLOMe. **(B)** Representative confocal images of STING and the trans-Golgi network marker TGN46 in WT and *Bltp3a*$^{KO}$ BPPNM cells treated with vehicle or LLOMe. DAPI marks nuclei. Data are shown as mean ± SEM. Each symbol represents an independent biological replicate. ns, not significant.

Figure S1

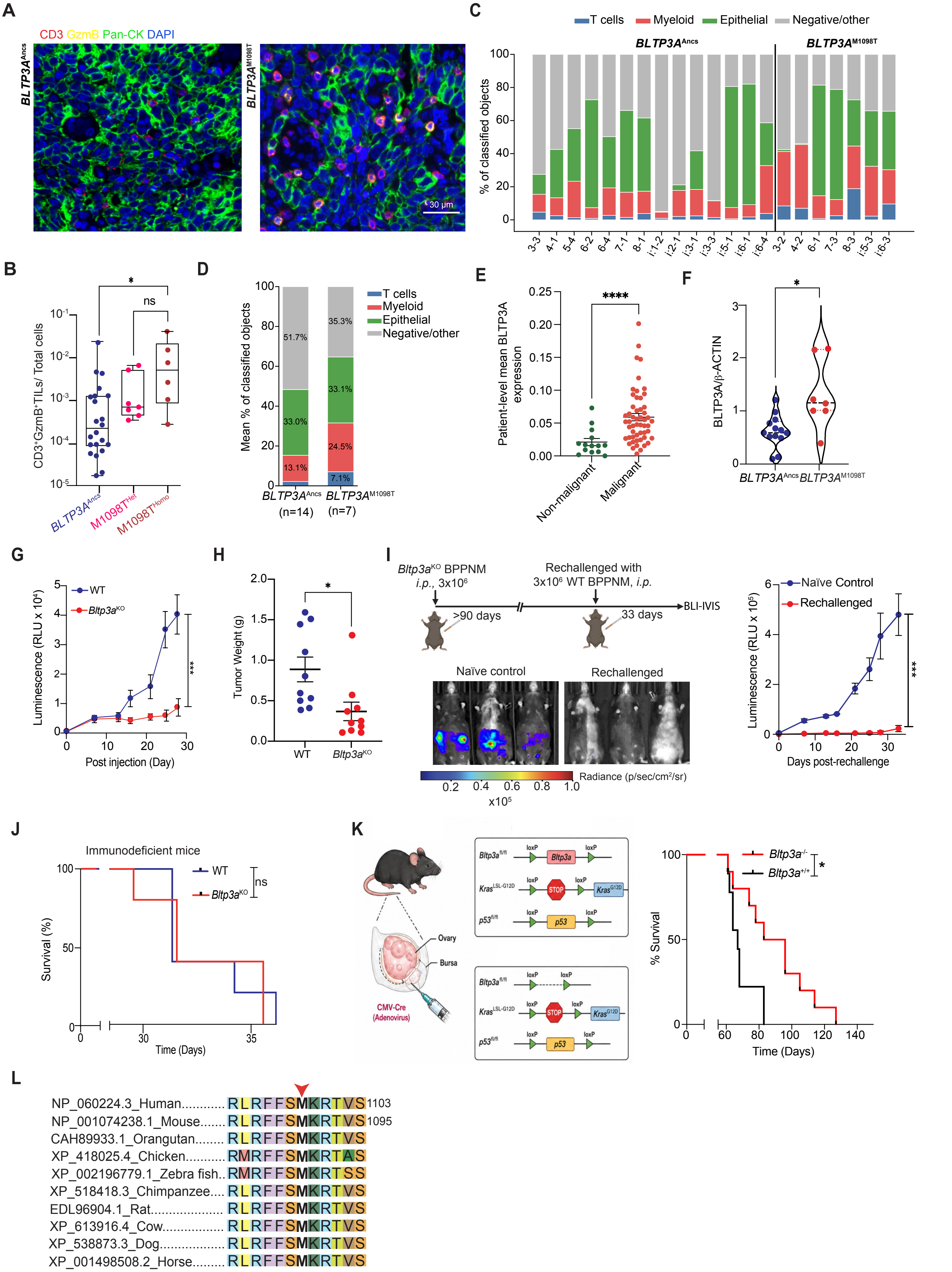

A
CD3 GzmB Pan-CK DAPI
BLTP3A^Ancs
BLTP3A^M1098T
30 µm
B
CD3⁺GzmB⁺TILs/ Total cells
*
ns
BLTP3A^Ancs
M1098T^Het
M1098T^Homo
C
T cells
Myeloid
Epithelial
Negative/other
BLTP3A^Ancs
BLTP3A^M1098T
% of classified objects
3-3
4-1
5-4
6-2
6-4
7-1
8-1
i:1-2
i:2-1
i:3-1
i:3-3
i:5-1
i:6-1
i:6-4
3-2
4-2
6-1
7-3
8-3
i:5-3
i:6-3
D
Mean % of classified objects
51.7%
33.0%
13.1%
35.3%
33.1%
24.5%
7.1%
BLTP3A^Ancs (n=14)
BLTP3A^M1098T (n=7)
E
Patient-level mean BLTP3A expression
****
Non-malignant
Malignant
F
BLTP3A/β-ACTIN
*
BLTP3A^Ancs
BLTP3A^M1098T
G
WT
Bltp3a^KO
Luminescence (RLU x 10⁴)
***
Post injection (Day)
H
Tumor Weight (g)
*
WT
Bltp3a^KO
I
Bltp3a^KO BPPNM
i.p., 3x10⁶
>90 days
Rechallenged with
3x10⁶ WT BPPNM, i.p.
33 days
BLI-IVIS
Naïve control
Rechallenged
Radiance (p/sec/cm²/sr)
x10⁵
Naïve Control
Rechallenged
Luminescence (RLU x 10⁵)
***
Days post-rechallenge
J
Immunodeficient mice
WT
Bltp3a^KO
ns
Survival (%)
Time (Days)
K
Ovary
Bursa
CMV-Cre (Adenovirus)
Bltp3a^fl/fl
Kras^LSL-G12D
p53^fl/fl
loxP
STOP
Kras^G12D
p53
Bltp3a^-/-
Bltp3a^+/+
*
% Survival
Time (Days)
L
NP_060224.3_Human............ RLRFFSMKRTVS 1103
NP_001074238.1_Mouse........ RLRFFSMKRTVS 1095
CAH89933.1_Orangutan........ RLRFFSMKRTVS
XP_418025.4_Chicken........... RMRFFSMKRTAS
XP_002196779.1_Zebra fish.. RMRFFSMKRTSS
XP_518418.3_Chimpanzee.... RLRFFSMKRTVS
EDL96904.1_Rat................... RLRFFSMKRTVS
XP_613916.4_Cow................. RLRFFSMKRTVS
XP_538873.3_Dog.................. RLRFFSMKRTVS
XP_001498508.2_Horse......... RLRFFSMKRTVS

### Figure S2

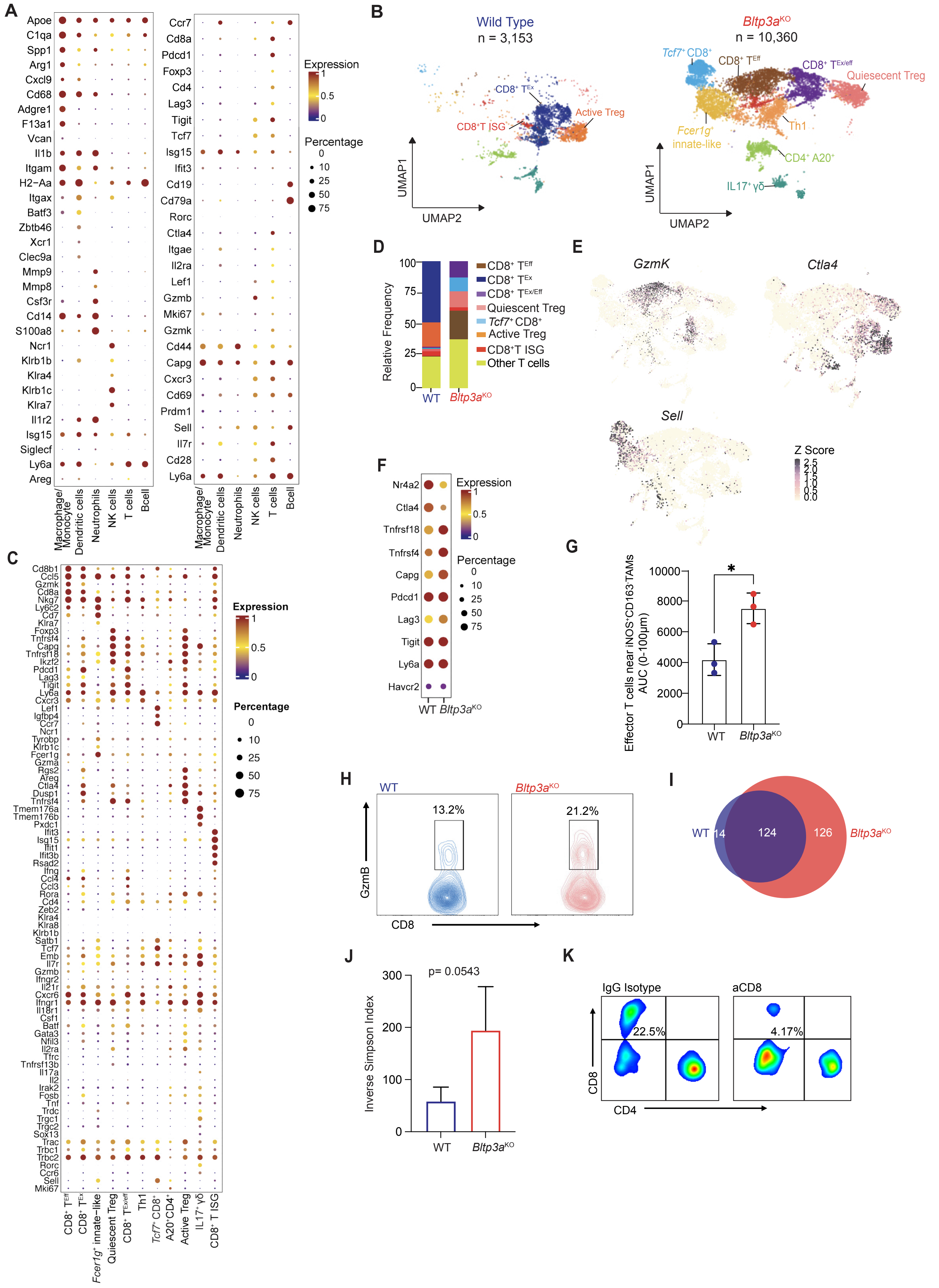
A
B
Wild Type
n = 3,153
Bltp3aKO
n = 10,360
UMAP1
UMAP2
C
D
Relative Frequency
CD8+ TEff
CD8+ TEx
CD8+ TEx/Eff
Quiescent Treg
Tcf7+ CD8+
Active Treg
CD8+T ISG
Other T cells
E
GzmK
Ctla4
Sell
Z Score
F
Expression
Percentage
G
Effector T cells near iNOS+CD163-TAMs AUC (0-100μm)
H
13.2%
21.2%
GzmB
CD8
I
WT
14
124
126
J
Inverse Simpson Index
p= 0.0543
K
IgG Isotype
aCD8
22.5%
4.17%
CD4

### Figure S3

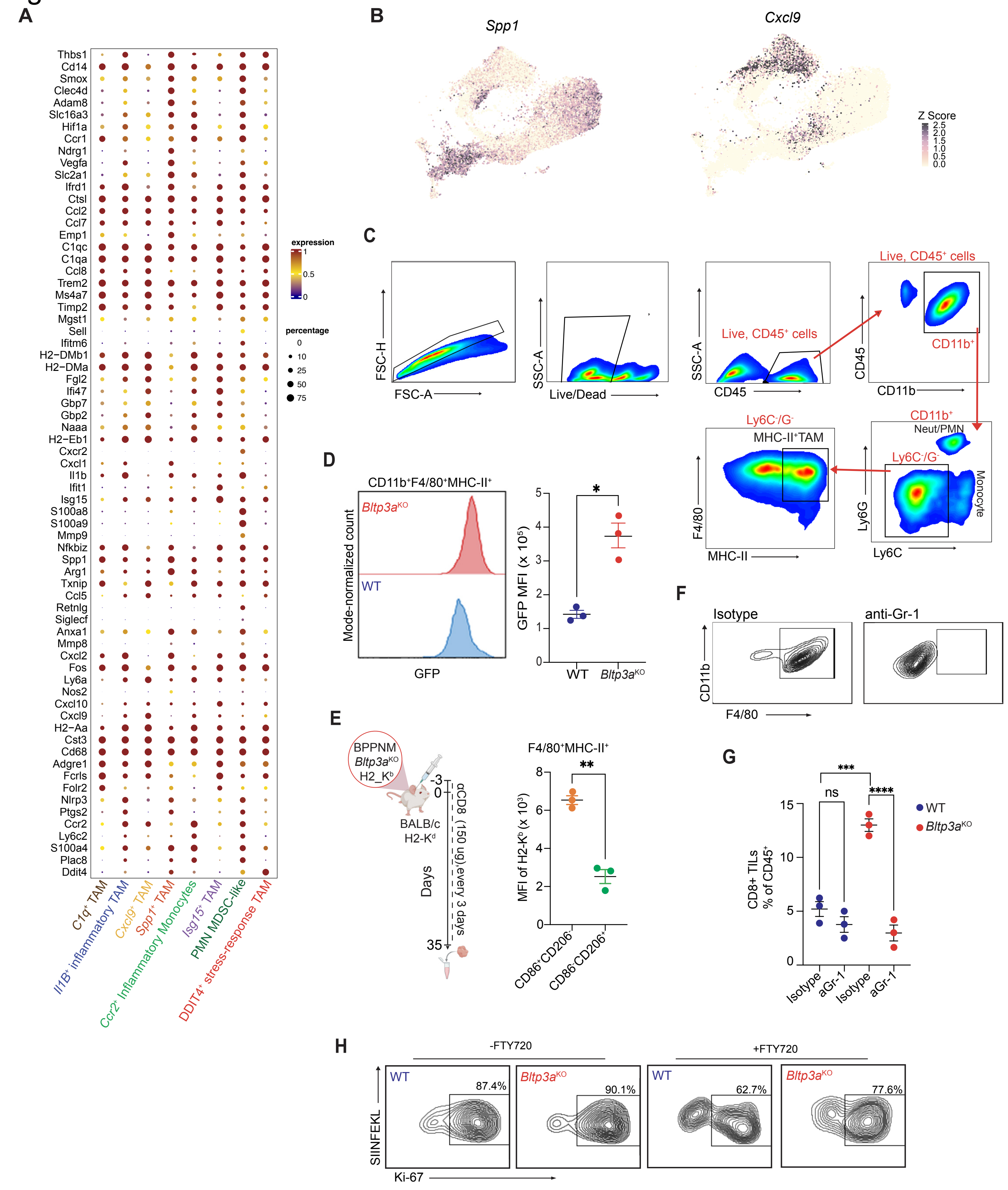

A
Thbs1
Cd14
Smox
Clec4d
Adam8
Slc16a3
Hif1a
Ccr1
Ndrg1
Vegfa
Slc2a1
Ifrd1
Ctsl
Ccl2
Ccl7
Emp1
C1qc
C1qa
Ccl8
Trem2
Ms4a7
Timp2
Mgst1
Sell
Ifitm6
H2−DMb1
H2−DMa
Fgl2
Ifi47
Gbp7
Gbp2
Naaa
H2−Eb1
Cxcr2
Cxcl1
Il1b
Ifit1
Isg15
S100a8
S100a9
Mmp9
Nfkbiz
Spp1
Arg1
Txnip
Ccl5
Retnlg
Siglecf
Anxa1
Mmp8
Cxcl2
Fos
Ly6a
Nos2
Cxcl10
Cxcl9
H2−Aa
Cst3
Cd68
Adgre1
Fcrls
Folr2
Nlrp3
Ptgs2
Ccr2
Ly6c2
S100a4
Plac8
Ddit4
expression
1
0.5
0
percentage
0
10
25
50
75
C1q+ TAM
Il1B+ inflammatory TAM
Cxcl9+ TAM
Spp1+ TAM
Ccr2+ Inflammatory Monocytes
Isg15+ TAM
PMN MDSC-like
DDIT4+ stress-response TAM
B
Spp1
Cxcl9
Z Score
2.5
2.0
1.5
1.0
0.5
0.0
C
FSC-H
FSC-A
SSC-A
Live/Dead
SSC-A
CD45
Live, CD45+ cells
Live, CD45+ cells
CD45
CD11b
CD11b+
CD11b+
Neut/PMN
Ly6C-/G-
Monocyte
Ly6G
Ly6C
Ly6C-/G-
MHC-II+TAM
F4/80
MHC-II
D
CD11b+F4/80+MHC-II+
Bltp3aKO
WT
Mode-normalized count
GFP
*
GFP MFI (x 10^5)
5
4
3
2
1
0
WT
Bltp3aKO
E
BPPNM
Bltp3aKO
H2_Kb
BALB/c
H2-Kd
-3
0
αCD8 (150 ug),every 3 days
Days
35
F4/80+MHC-II+
**
MFI of H2-Kb (x 10^3)
8
6
4
2
0
CD86+CD206-
CD86-CD206+
F
Isotype
anti-Gr-1
CD11b
F4/80
G
***
ns
****
WT
Bltp3aKO
CD8+ TILs
% of CD45+
15
10
5
0
Isotype
aGr-1
Isotype
aGr-1
H
-FTY720
+FTY720
WT
87.4%
Bltp3aKO
90.1%
WT
62.7%
Bltp3aKO
77.6%
SIINFEKL
Ki-67

### Figure S4

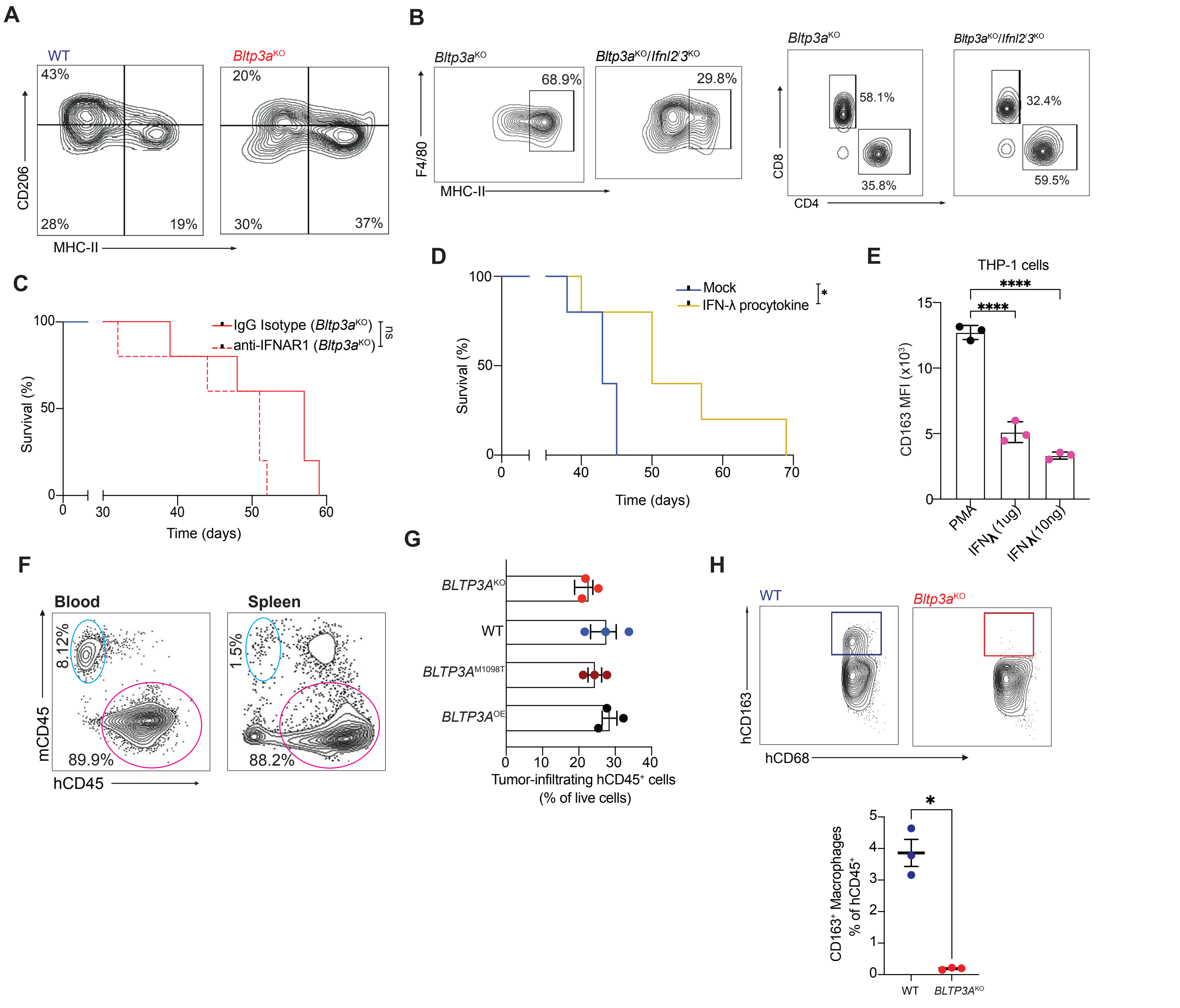

A
WT
Bltp3aKO
43%
28%
19%
20%
30%
37%
CD206
MHC-II
B
Bltp3aKO
Bltp3aKO/Ifnl2/3KO
68.9%
29.8%
F4/80
MHC-II
58.1%
35.8%
32.4%
59.5%
CD8
CD4
C
IgG Isotype (Bltp3aKO)
anti-IFNAR1 (Bltp3aKO)
ns
Survival (%)
Time (days)
D
Mock
IFN-λ procytokine
*
Survival (%)
Time (days)
E
THP-1 cells
****
****
CD163 MFI (x10^3)
PMA
IFNλ(1ug)
IFNλ(10ng)
F
Blood
Spleen
8.12%
89.9%
1.5%
88.2%
mCD45
hCD45
G
BLTP3AKO
WT
BLTP3AM1098T
BLTP3AOE
Tumor-infiltrating hCD45+ cells
(% of live cells)
H
WT
Bltp3aKO
hCD163
hCD68
*
CD163+ Macrophages
% of hCD45+
WT
BLTP3AKO

### Figure S5

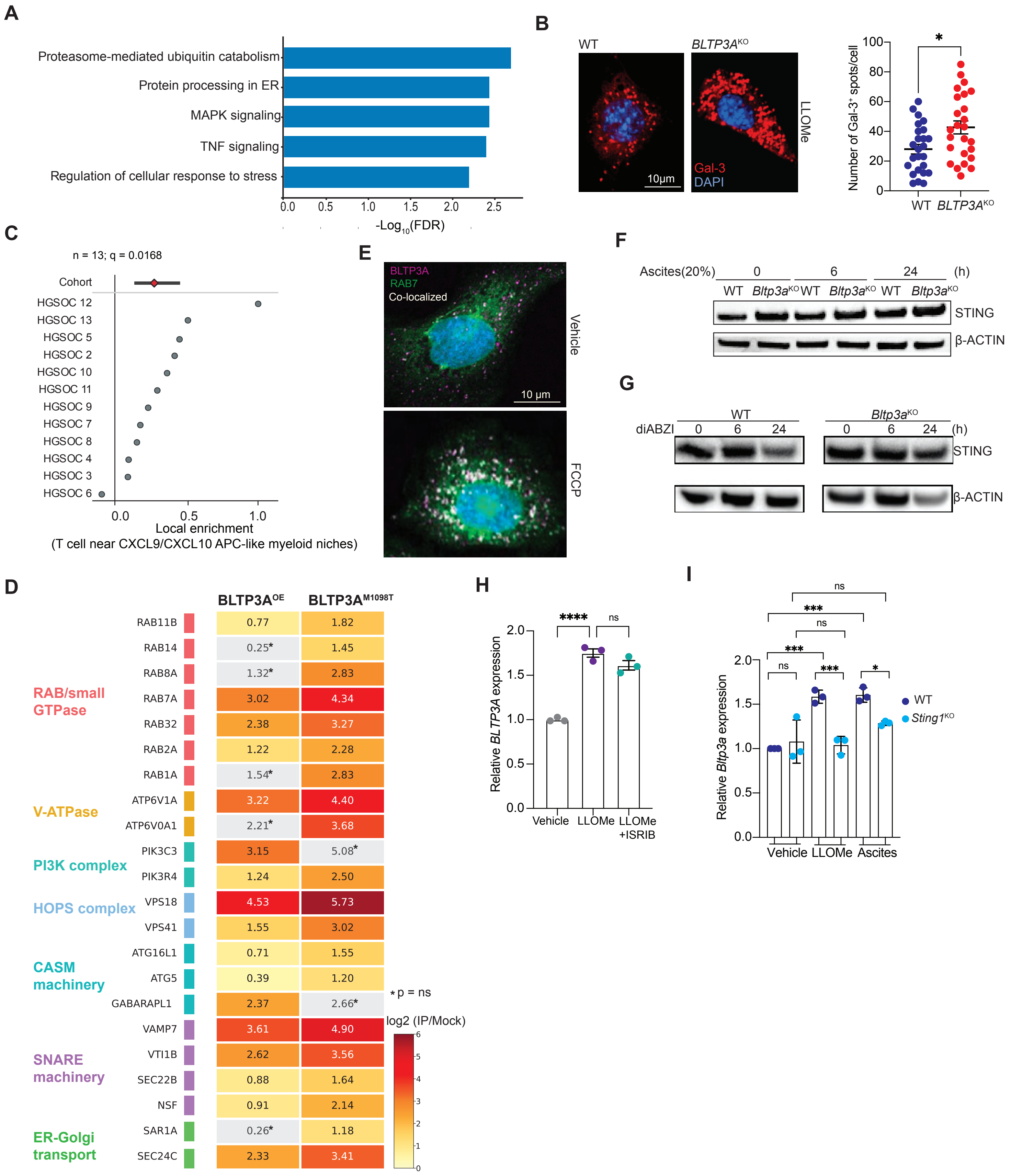
A
Proteasome-mediated ubiquitin catabolism
Protein processing in ER
MAPK signaling
TNF signaling
Regulation of cellular response to stress
0.0 0.5 1.0 1.5 2.0 2.5
-Log10(FDR)
B
WT
BLTP3A^KO
LLOMe
10μm
Gal-3
DAPI
Number of Gal-3+ spots/cell
0 20 40 60 80 100
*
WT BLTP3A^KO
C
n = 13; q = 0.0168
Cohort
HGSOC 12
HGSOC 13
HGSOC 5
HGSOC 2
HGSOC 10
HGSOC 11
HGSOC 9
HGSOC 7
HGSOC 8
HGSOC 4
HGSOC 3
HGSOC 6
0.0 0.5 1.0
Local enrichment
(T cell near CXCL9/CXCL10 APC-like myeloid niches)
E
BLTP3A
RAB7
Co-localized
Vehicle
10 μm
FCCP
F
Ascites(20%) 0 6 24 (h)
WT Bltp3a^KO WT Bltp3a^KO WT Bltp3a^KO
STING
β-ACTIN
G
diABZI
WT
Bltp3a^KO
0 6 24 (h)
0 6 24
STING
β-ACTIN
D
BLTP3A^OE
BLTP3A^M1098T
RAB/small GTPase
RAB11B 0.77 1.82
RAB14 0.25* 1.45
RAB8A 1.32* 2.83
RAB7A 3.02 4.34
RAB32 2.38 3.27
RAB2A 1.22 2.28
RAB1A 1.54* 2.83
V-ATPase
ATP6V1A 3.22 4.40
ATP6V0A1 2.21* 3.68
PI3K complex
PIK3C3 3.15 5.08*
PIK3R4 1.24 2.50
HOPS complex
VPS18 4.53 5.73
VPS41 1.55 3.02
CASM machinery
ATG16L1 0.71 1.55
ATG5 0.39 1.20
GABARAPL1 2.37 2.66*
SNARE machinery
VAMP7 3.61 4.90
VTI1B 2.62 3.56
SEC22B 0.88 1.64
NSF 0.91 2.14
ER-Golgi transport
SAR1A 0.26* 1.18
SEC24C 2.33 3.41
* p = ns
log2 (IP/Mock)
0 1 2 3 4 5 6
H
Relative BLTP3A expression
0.0 0.5 1.0 1.5 2.0
****
ns
Vehicle LLOMe LLOMe +ISRIB
I
Relative Bltp3a expression
0.0 0.5 1.0 1.5 2.0
ns
***
ns
***
ns
***
*
WT
Sting1^KO
Vehicle LLOMe Ascites

### Figure S6

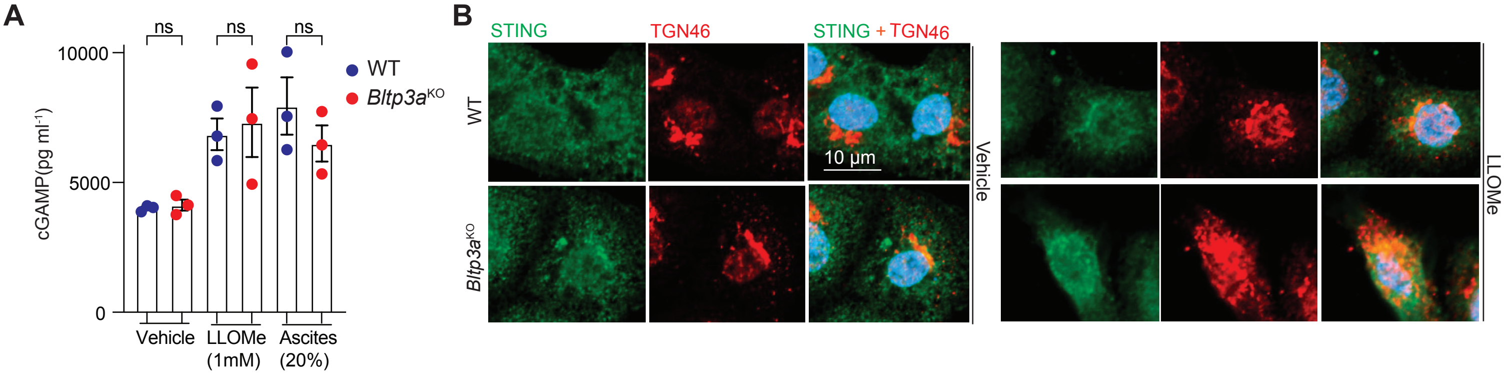

A
ns
ns
ns
10000
5000
0
cGAMP(pg ml-1)
WT
Bltp3aKO
Vehicle
LLOMe (1mM)
Ascites (20%)
B
STING
TGN46
STING + TGN46
WT
Bltp3aKO
10 μm
Vehicle
LLOMe
